# Transcriptomic study of WSSV infection in *Litopenaeus vannamei* lymphoid organ via single nuclei RNA sequencing

**DOI:** 10.64898/2025.12.09.693182

**Authors:** Alexandra Florea, Nick Wade, Sarah J. Salisbury, James Furniss, Clémence Fraslin, Ambre Chapuis, Kallen Sullivan, Robert Stewart, Diego Robledo, Tim P. Bean

## Abstract

**Background:** Aquaculture is the fastest growing farmed food sector and is a key part of global food security. Crustacean aquaculture, while being one of the most profitable sectors, is threatened by pathogenic diseases such as white spot disease (WSD), which results in severe stock losses and threatens animal health and welfare. In this study, we used novel techniques to study the impact of white spot syndrome virus (WSSV), the causative agent of WSD, on Pacific whiteleg shrimp (*Litopenaeus vannamei*), and uncover additional information that may be used in creating WSSV-resistant shrimp stocks.

**Results:** We successfully developed a novel nuclei isolation protocol optimized for shrimp tissues to prepare our samples for single nuclei RNA-sequencing following a pathogen challenge comprising of 32 adult whiteleg shrimp infected with WSSV either through their feed or by injection. We constructed the first penaeid shrimp lymphoid organ cell atlas to improve our understanding of the characteristics of this immune organ, constructed the UMAPs and identified marker genes that define distinct cell clusters to reveal the biological functions of different cell types within this organ.

**Conclusion:** By comparing gene expression between the control and WSSV-infected samples we uncovered multiple genes of interest that that have the potential to be used as targets for CRISPR gene editing for WSSV-resistance in Pacific whiteleg shrimp. These genes have the potential to be valuable assets for future gene editing studies in commercially important penaeid shrimp species.

**Graphical Abstract:** 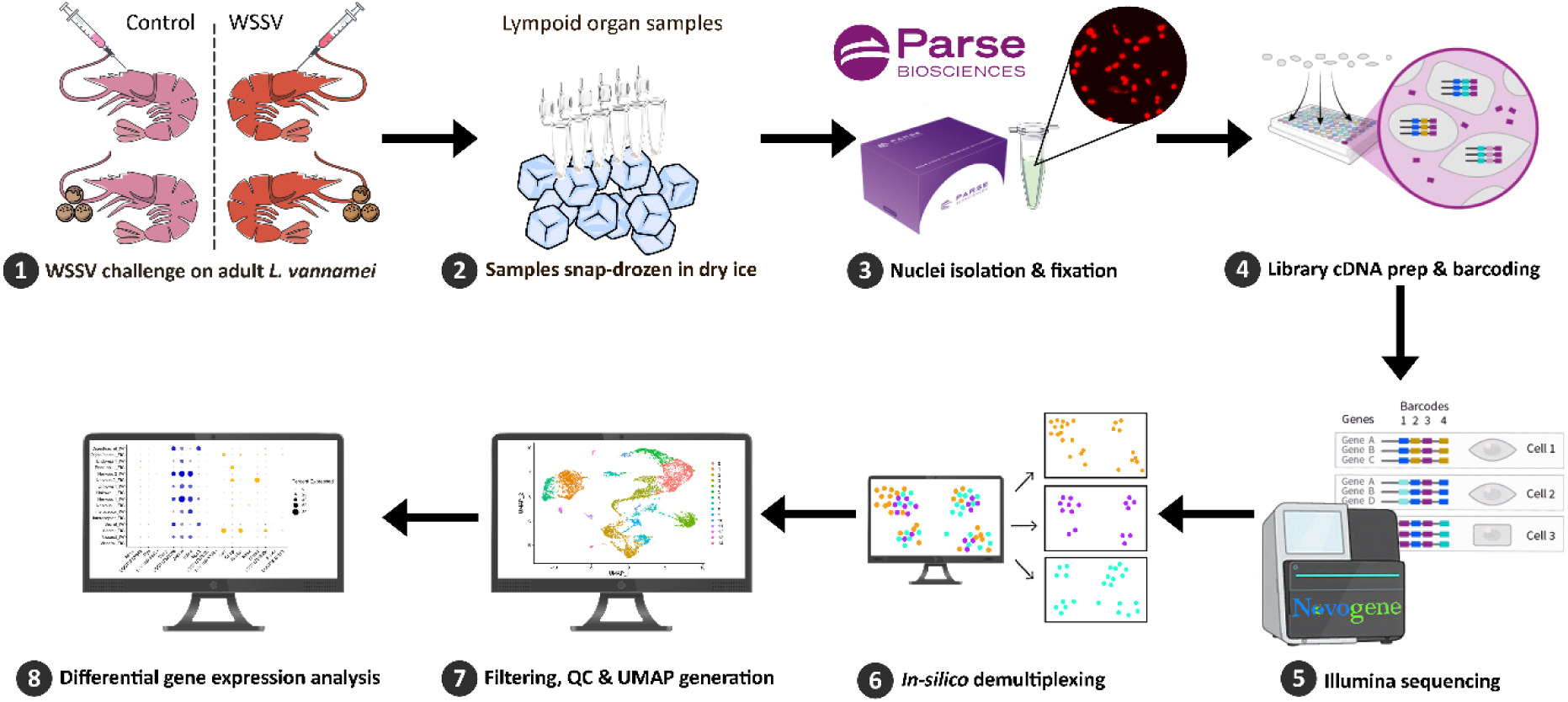

## 1. Introduction

Aquaculture contributes significantly to global food production, and its importance is projected to rise. By 2032, aquaculture is estimated to account for 54 percent of the total production of aquatic animals (1). Penaeid shrimp, such as Pacific whiteleg shrimp (*Litopenaeus vannamei*), Chinese white shrimp (*Fenneropenaeus chinensis*), kuruma shrimp (*Marsupenaeus japonicus*) and black tiger shrimp (*Panaeus monodon*), accounted for 62.2% of all farmed crustacean species, and remain the second most valuable farmed group after salmonids, accounting for 17% of total global aquaculture value (1). Whiteleg shrimp was the top produced aquaculture species globally, at 6.8 million tonnes.

Despite their global productivity, shrimp farms frequently face disease outbreaks which severely hamper their production (2). White spot syndrome virus (WSSV) is recognised as one of the most significant threats to crustacean aquaculture by the World Organization of Animal Health (3), and effective disease control methods are urgently needed.

Invertebrate immune responses are less well characterised than those of mammals, primarily due to their evolutionary distance from humans and mice, where the majority of functional biology studies have been focused (4). Unlike mammals, penaeid shrimp lack developed antibody-mediated adaptive immunity, relying solely on effective cellular and humoral innate immunity, which in turns limits the potential for vaccine development (5); (6). In addition, crustacean immunity and the interactions between WSSV and host shrimp cells are not well characterised, therefore limiting the development of effective control measures.

It is known that the hepatopancreas, lymphoid organ and haemocytes play crucial roles in controlling the innate immunity responses in penaeid shrimp, making it essential to study their response to pathogens to facilitate disease prevention and resource development for future research (7). The lymphoid organ, also known as the Oka organ, plays a major immunological role alongside haemolymph cells and hepatopancreas. It is a paired, lobed, immune organ found in penaeid shrimp, and other crustacean species such as *Macrobrachium rosenbergii* (8) and *Procambarus clarkii* (9). It functions in pathogen defence by facilitating the recognition and elimination of viral and bacterial invaders, as evidenced by its significant role during infections by pathogens such as Vibrio parahaemolyticus and WSSV. Mechanisms by which the lymphoid organ fights against invading pathogens are believed to include phagocytosis, filtering of foreign materials from the haemolymph, bacteriostasis and viral degradation via the lymphoid organ spheroid (LOS) cells (10); (7); (11).

Transcriptomic analyses have shown differential gene expression in the lymphoid organ in response to various pathogens, indicating its adaptive immune functions. For instance, Vibrio infections trigger broad immune responses, including the activation of pattern recognition receptors and phagocytosis-related genes, whereas WSSV infections tend to suppress immune responses, highlighting the organ’s dynamic role in managing shrimp immunity against diverse threats (11).

Given the potential immunological roles of multiple cell types, distinguishing between them is critical to understanding their individual and collective contributions to a host’s immune response. For instance, it is estimated that humans have almost 200 different cell types, many of which are well documented (12). In contrast, few cell types have been documented in crustaceans and they generally lack thoroughly characterised marker genes, or cluster of cellular differentiation marker genes (CD markers) (6). Consistent cell type characterizations have been elusive due to variations in cell classification, study methodologies, and the absence of genetic and annotation tools for shrimp cell populations. Studying crustacean immunity at the cellular level would provide insights into the evolution of immune mechanisms and identify fundamental immune processes that are conserved across species. This approach also provides the opportunity to uncover novel immune strategies, improve disease models, and advance biotechnological applications and drug discovery.

Advances in next-generation sequencing technologies, particularly single-cell (scRNA-seq) and single nuclei RNA sequencing (snRNA-seq), have enabled comprehensive analysis of mRNA at the individual cell level, facilitating the study of gene expression differences among single cells (13). To date sc/snRNA-seq has been applied to a variety of aquatic animal species including Atlantic salmon, rainbow trout, turbot, grouper, orange spotted grouper, Nile tilapia, Asian sea bass, Kuruma shrimp, white shrimp, black tiger shrimp and Hong Kong Oyster, revealing cellular diversity and gene expression patterns (summarized in (14)). Beyond identifying and transcriptionally characterizing cell types, snRNA-seq has also proven valuable in revealing the specific cell types responding to pathogens as well as those cell types targeted by pathogens to modulate their host’s immune response (e.g., (15)).

Capitalizing on the strengths of this technology, snRNA-seq studies on penaeid shrimp species have started to emerge across the literature, focusing on haemocyte and hepatopancreas tissues, studying cell types and differentiation (16); (17), and differentially expressed genes (DEGs) during events such as ammonia nitrogen stress (18), cold stress (19), and artificial infections via injection with WSSV inoculum (6); (13) (**Table 1**).

**Table 1.**
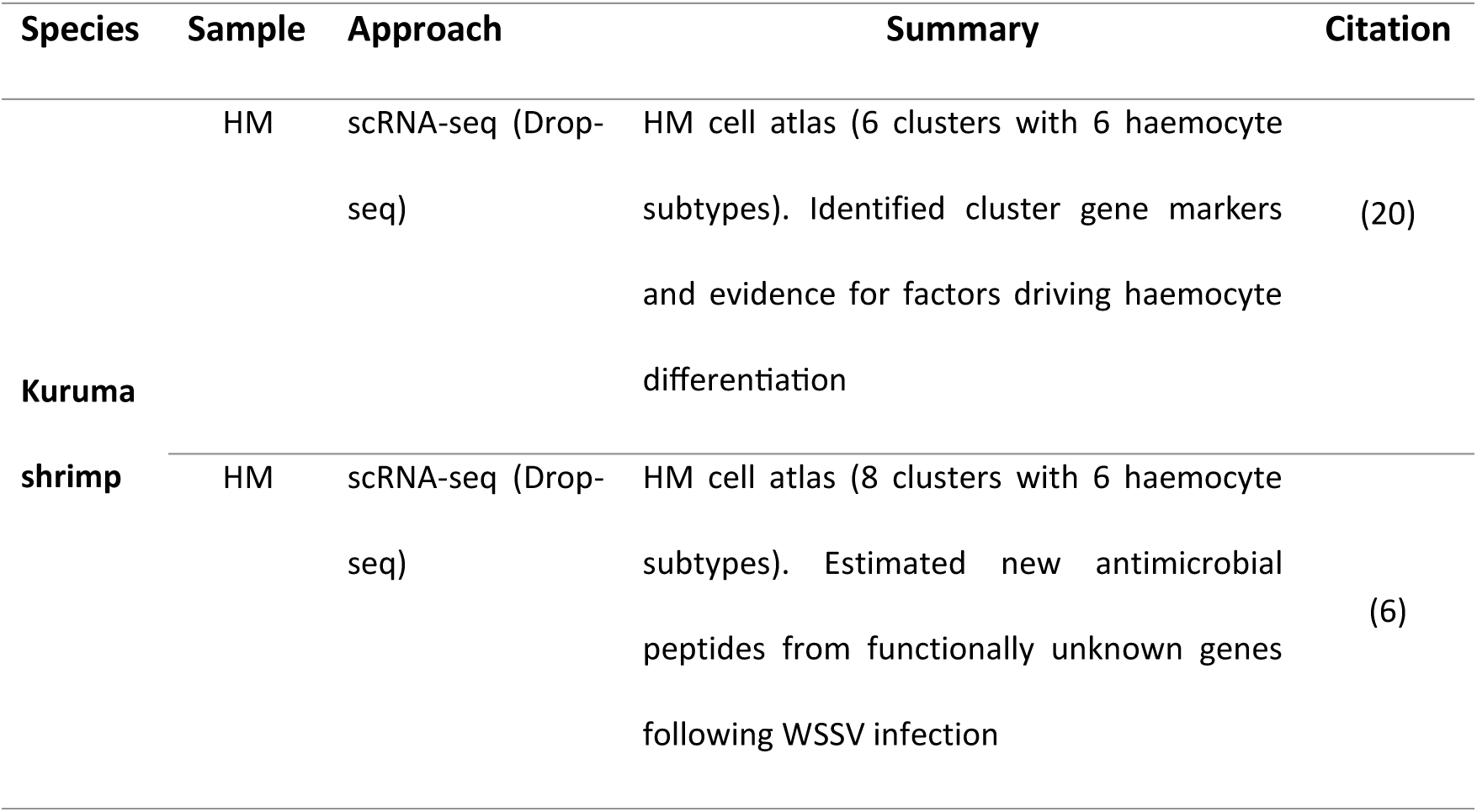
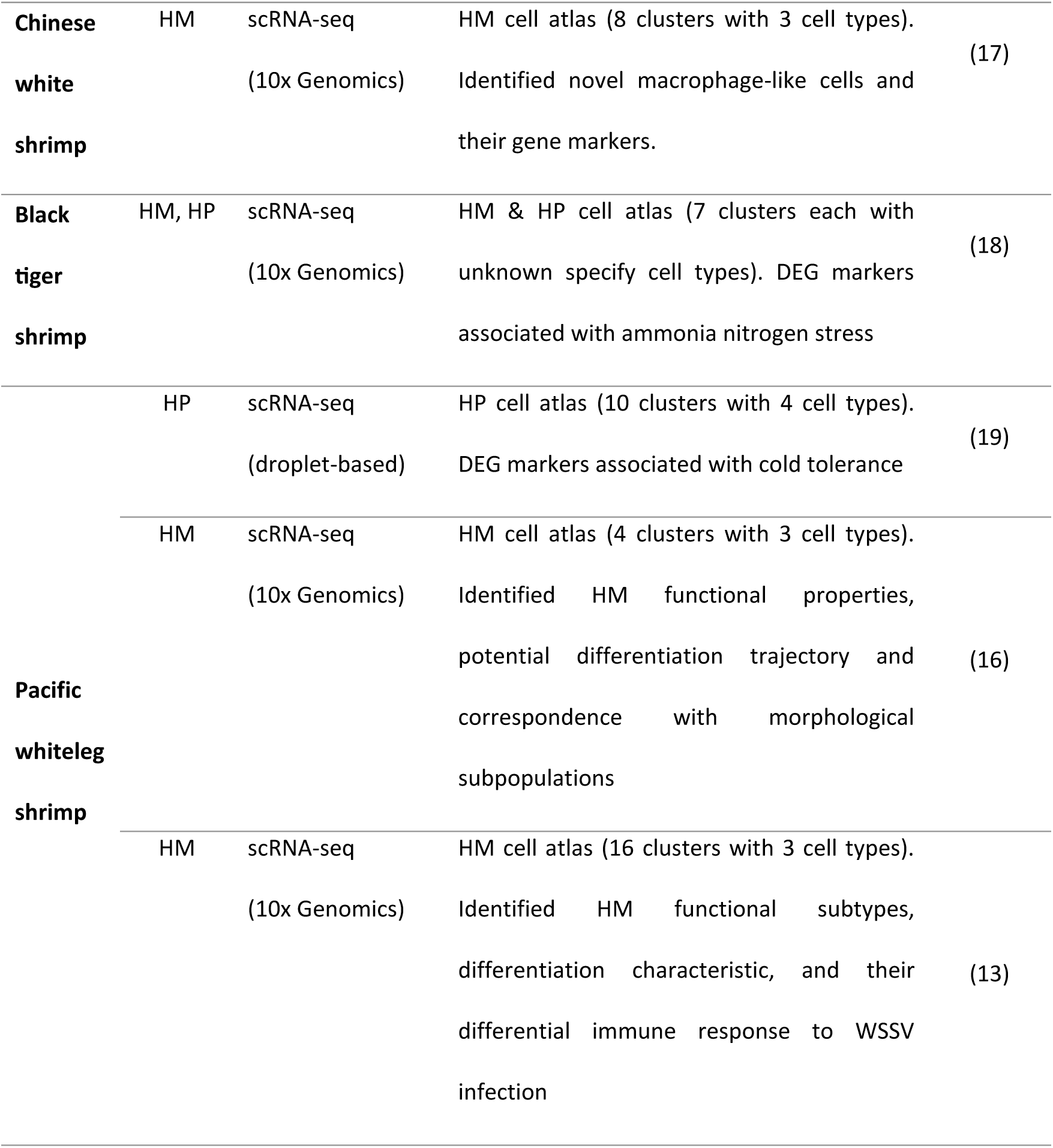
Summary of single cell and single nuclei RNA-sequencing studies in penaeid shrimp of aquaculture importance.

In this study, we performed snRNA sequencing on lymphoid organ tissue of healthy and WSSV-infected *L. vannamei,* with the aim of identifying the different cell types found in the organ, including both functional and structural-type cells, with particular interest to immune-tissue cell populations and sub-populations. At a gene-level, we analysed differentially expressed genes pre- and post-infection and singled-out several genes related to viral immunity.

## 2. Materials and methods

### 2.1 Experimental animal care

Adult *L. vannamei* weighting 6-12 g were used for this study and held in the Recirculating Aquacultur System (RAS) shrimp facility at the Roslin Institute (Midlothian, UK) (**Figure 1**). The shrimp were kept at 28 ± 0.5 °C, in 3 L or 10 L plastic tanks containing aerated artificial seawater with a salinity of 25‰ and pH of 8.0. The shrimp were fed a commercial shrimp pellet diet twice a day (∼5% body weight/day). The shrimp were housed in individual tanks and solid waste syphoning, and water tests were performed daily. The aquaria connected to the RAS system had an average water turnover of 3L/h which was filtered using mechanical and biological filters before being passed through a protein skimmer and returned to the system.

**Figure 1.**
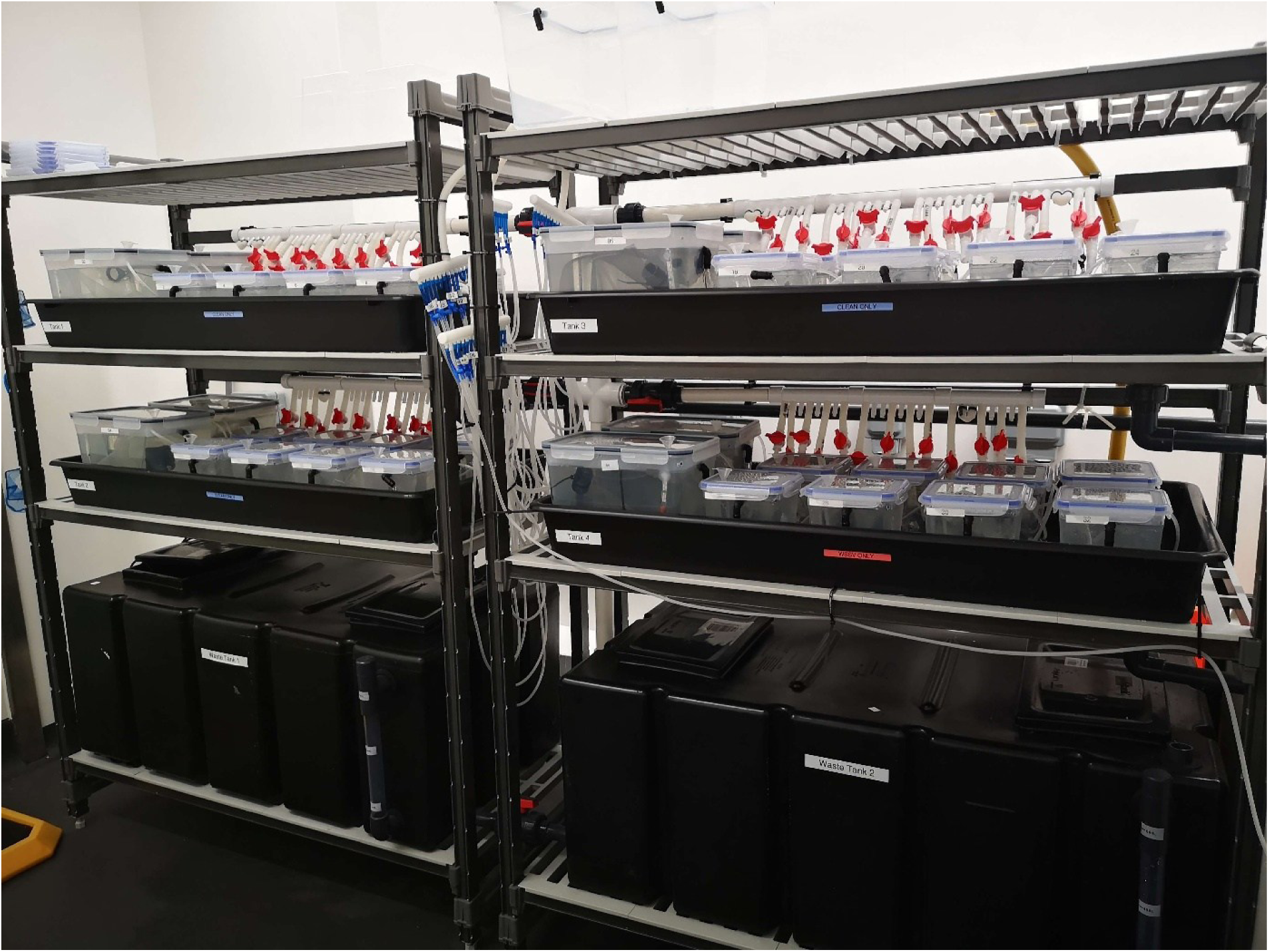
Shrimp holding facility. The picture shows the RAS system setup used for the viral challenges in adult L. vannamei shrimp. Each tank is fitted with an individual aeration system. A small-scale RAS system is used to keep the water parameters constant and water quality within safety levels. The feeding was done manually.

### 2.2 WSSV quantification

A WSSV-standard curve was used to quantify all WSSV samples. The qPCRs were done with Brilliant III SYBR Master Mix (Agilent), with published primer and probe sequences (21) (**Table 2**). The reaction volumes and the thermocycler program were prepared and run according to manufacturer’s instructions.

**Table 2.**
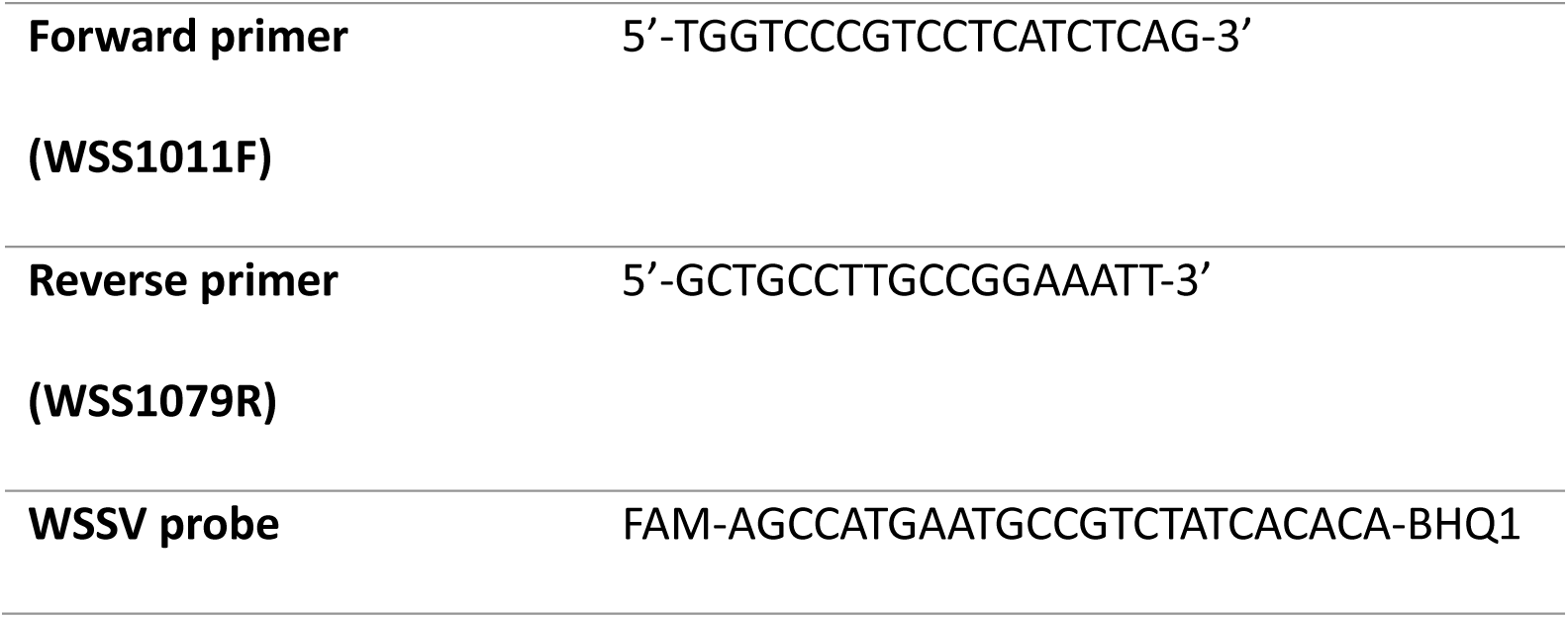
Primers and probe used for WSSV 69bp amplicon amplification in qPCR.

### 2.3 WSSV stock expansion

Purified viral stock and infected shrimp tissue were sourced from CEFAS - *Centre of Environment Fisheries and Aquaculture Science* (Lowestoft, England). We expanded our viral stock by bath challenge and feed challenges; each performed on adult *L. vannamei* shrimp. For the bath challenge we decanted 2 mL vial of purified WSSV stock into a 20 L aquarium with three adult shrimp. Separately, we prepared infected shrimp material and mixed it with the standard food pellets which was fed to six adult shrimp. The challenges were performed in two isolated aquaria with filtration and heating. Both groups were maintained for 7 days before shrimps were sacrificed on ice. The tail muscle of the sacrificed shrimps was extracted and macerated using a tissue homogenizer. All the tissue samples were tested for viral load using qPCR. Viral tissue stocks that exceeded 1.5 × 10^6^ WSSV copies/µl were kept frozen at −80 °C until their use in the main WSSV challenge.

### 2.4 WSSV infection and viral load quantification *in-vivo*

For our experimental viral challenge we used 32 adult shrimp, split into two main groups of 16 shrimp: control and infected. Each of the groups were further split in two eight-shrimp sub-groups by infection method: feed and injection. The WSSV-challenged shrimp were maintained in isolated aquaria which were closed-off from the recirculation system for the duration of the challenge in order to prevent cross-contamination of the control group. The aquaria on this tray had daily manual water exchanges, with the displaced water going straight into the waste tank for decontamination. To ensure identical conditions in both groups, the control shrimp were held in equivalent static systems with manual water changes.

Injection challenges were performed with 50 µl of saline solution (Tris-HCl 20mM & NaCl 400mM in dH_2_O, pH 7.4, 0.45 µm syringe filtered) delivered intra-muscularly in the dorsal side between the 3^rd^ and 4^th^ abdominal segment using a 26G needle. Animals in the WSSV-challenged group were injected with a mix of base saline solution and 5 µm-filtered WSSV stock, prepared from the previous challenge (approx. 1.5 × 10^6^ WSSV copies/µl) at a 50:1 ratio. The control group was injected using 100% saline solution.

For the feed challenge group, we prepared balls of food made out of macerated shrimp tail muscle tissue and shrimp food pellets at a 3:1 ratio. The balls were roughly 0.5 cm^3^. For the infected group, the muscle tissue came from the WSSV-infected shrimp (approx. 3 × 10^5^ WSSV copies/µl), while for the control group we used WSSV-free SPF shrimp (CEFAS, Lowestoft, England)

Post-infection, all shrimps were fed a normal diet twice a day. The WSSV-infected shrimp were monitored several times during the day, between early morning and later afternoon, for mortality and infection symptoms. Due to the infection timeline being slightly different for each group, the shrimp were sampled separately between 48 and 72h post-infection for the injection group (rapid infection, following an artificial infection method) and between 5- and 7-days post-infection for the feed group (slow infection, following a natural mechanism of infection) to ensure they were all in the same mid to late infection stage (**Figure 2**). Mortalities were discarded to avoid capturing cell death processes in transcriptomic analysis. Data on weight and moult stage was collected upon sacrificing the shrimp. Only inter-moult stage shrimp (C-D1) were used for the downstream analysis to ensure differential gene expression was not biased due to differences in moult stage.

**Figure 2.**
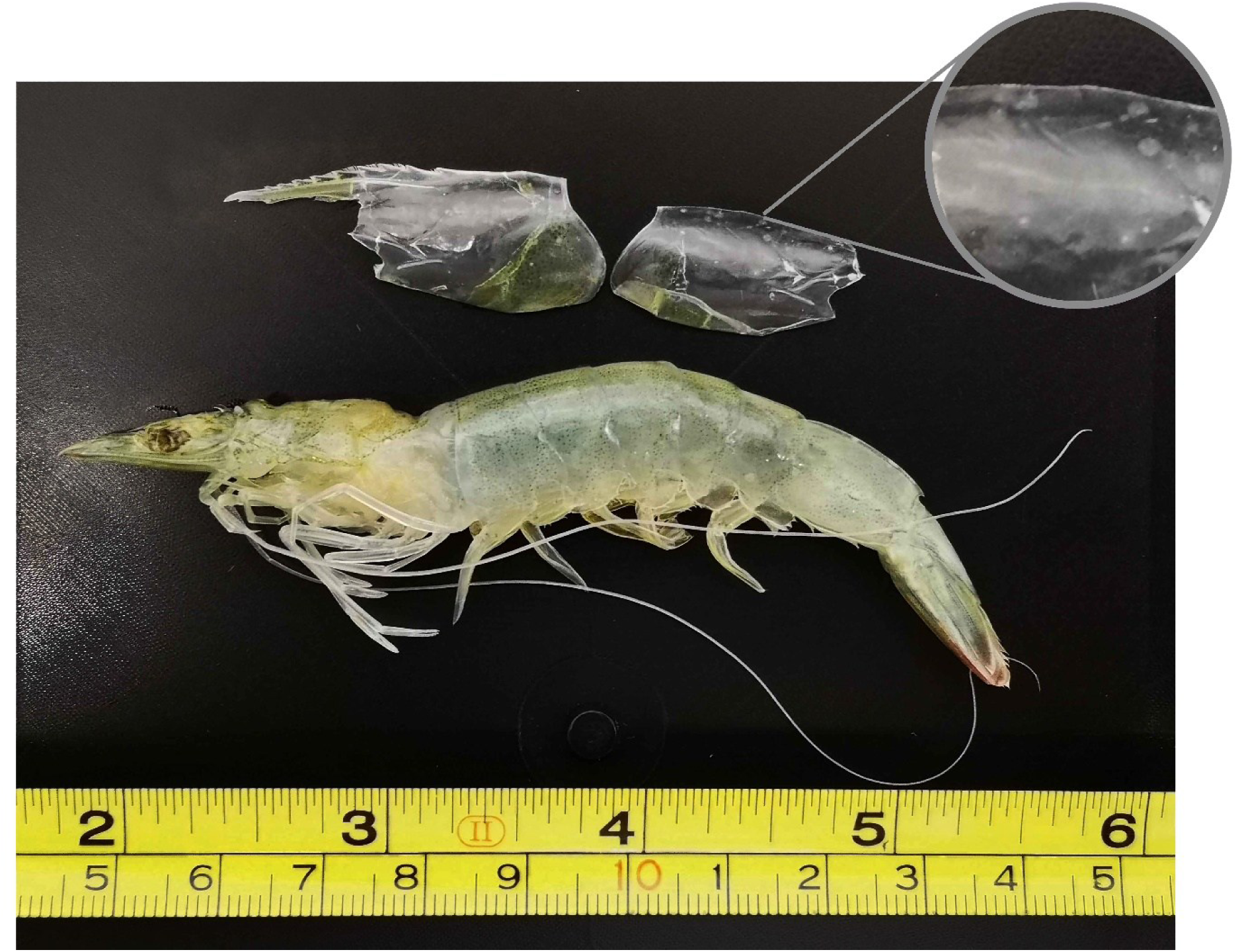
White spot disease symptoms in WSSV-challenged shrimp from the ‘feed’ group. The top-right close-up image shows white-spots on the carapace which are characteristic of white spot disease.

### 2.5 Sampling and nuclei processing

#### 2.5.1 Tissue sampling

Following the WSSV trial, all the shrimp with suitable moult stages were sacrificed on ice. Th individuals were dissected; the lymphoid organs were extracted (**Figure 3**) and then immediately frozen on dry ice. Tissues were stored at −80 ° C until processing.

**Figure 3.**
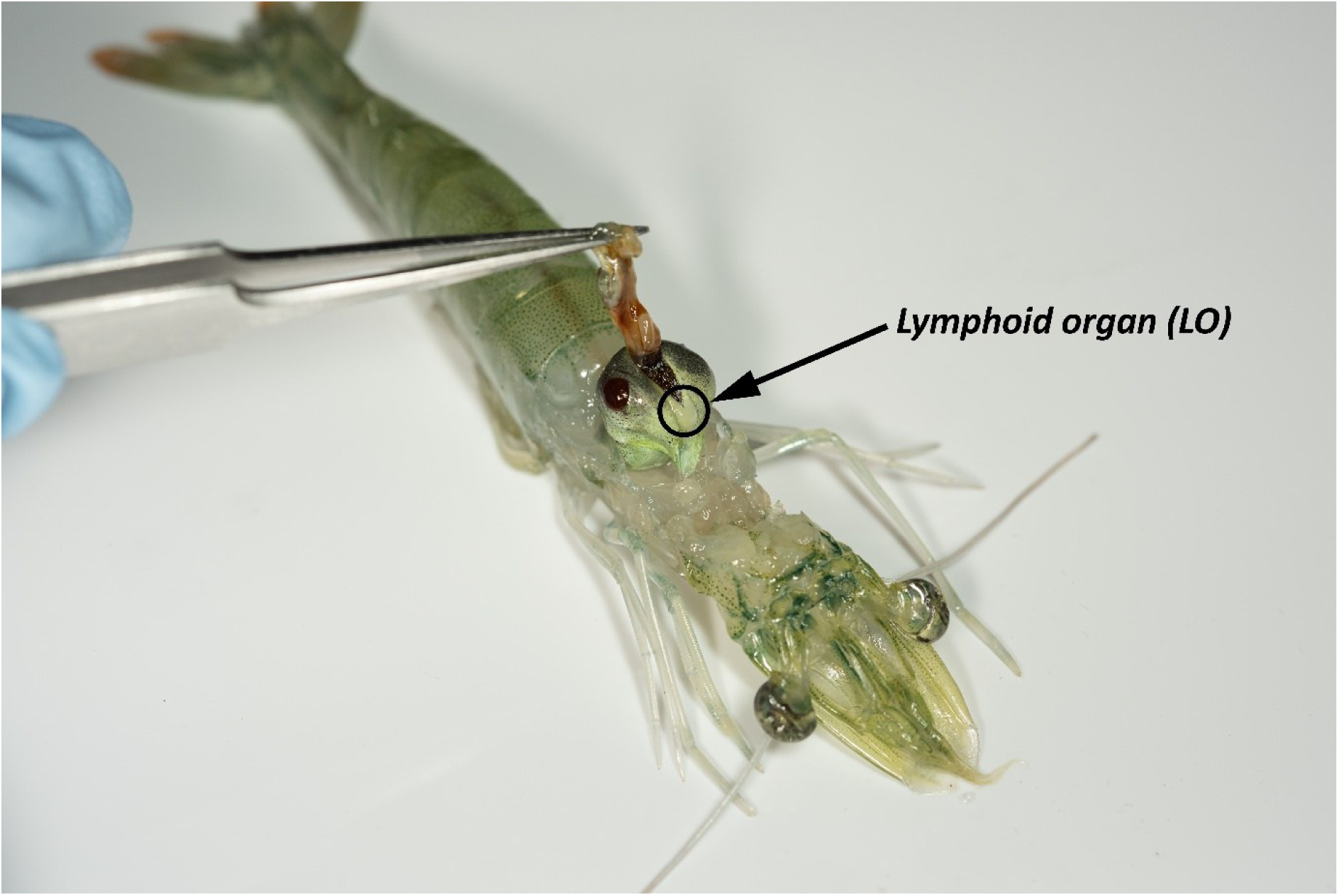
Pacific whiteleg shrimp dissection. The arrow and circle show lymphoid organ harvested for the transcriptomic analysis.

A muscle sample from each individual was subsequently tested for the presence or absence of WSSV (viral load). The 17 lymphoid organ pairs carried forward for the snRNA-seq experiment were chosen for their intermediate moult stage, absence of WSSV in control groups and high viral load in WSSV-positive samples

#### 2.5.2 Nuclei isolation and fixation

The nuclei were extracted using a modified version of the TST-based method (22). Briefly, we prepared the 2X stock of salt-Tris solution (ST buffer), 1X ST-buffer solution working solution (TST) as below:

- **2X ST buffer** (for 2 samples):

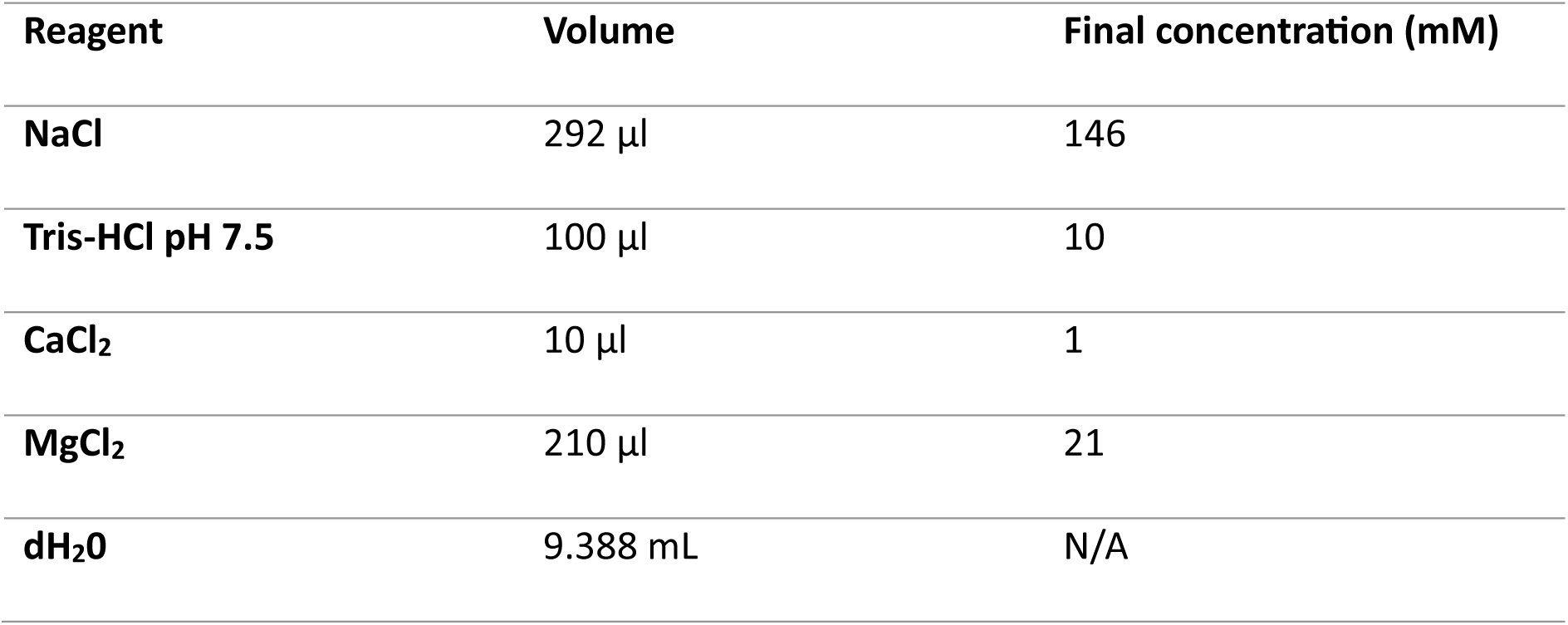
- **1X ST buffer:** Dilute 2x in nuclease-free water (1:1) and add 5.2 µl of RNAse Inhibitor (final conc. 40 U/mL).
- **TST solution:** Mix 2 mL of 2X ST buffer, 120 µl of Tween-20 1%, 20 µl BSA 2% and 1.86 mL of dH_2_0.

| Reagent | Volume | Final concentration (mM) |
| --- | --- | --- |
| NaCl | 292 µl | 146 |
| Tris-HCl pH 7.5 | 100 µl | 10 |
| CaCl<sub>2</sub> | 10 µl | 1 |
| MgCl<sub>2</sub> | 210 µl | 21 |
| dH<sub>2</sub>O | 9.388 mL | N/A |

For each sample, a 6-well tissue culture plate was placed on ice and 1 mL of TST buffer was added to a well. The frozen tissue sample (∼10-20 mg) was placed in the buffer straight from dry ice. Note that due to the lymphoid organs being very small, we could only get sufficient material to process with the Parse kit by pooling samples together. Lymphoid organ pairs from 2-3 individuals were pooled together for each sample depending on their size. The sample was then macerated as finely as possible with spring scissors for 4 minutes. The sample was then sequentially pipetted for 30 seconds with a P1000 and 30 seconds with a P200. Each sample lysate was then filtered through a 40 μm Falcon cell-strainer into a clean well of the plate, and then again through a 20 μm cell strainer. 1 mL of TST was added to wash the initial maceration well and material passed through the filters again to collect the remaining nuclei. The volume was brought up to 5 mL with the addition of 3 mL of 1X ST buffer which also inactivates the Tween-20. The lysate was transferred to a 15 mL Falcon tube and centrifuged at 4 °C for 5 minutes at 500 g. The pellet was re-suspended in between 200 µl 1X ST buffer.

A 9 µl sub-sample of the nuclei was taken and stained with 1 µl of Acridine Orange/Propidium Iodide. The LUNA-FX7™ Automated Cell Counter was used to count the nuclei as well as to determine the nuclei quality, degradation level, shape, size, and non-singlet percentage.

Following nuclei isolation and removal of a sub-sample for quality testing, nuclei were fixed immediately using the Evercode™ Nuclei Fixation v2 kit from Parse Biosciences according to manufacturer’s instructions. At the end, the nuclei were re-counted as above using the LUNA-FX7™ Automated Cell Counter. Finally, the fixed nuclei samples were chilled using a Mr. Frosty™ freezing container and stored at −80 °C until all other samples had been processed.

### 2.6 RNA library generation and sequencing

Once all the nuclei had been processed, approximately 5000 nuclei from each sample were prepared for sequencing using the Evercode™ WT Mega v2 from Parse Biosciences according to manufacturer’s instructions. Six total libraries were prepared using the 17 lymphoid organ pairs as follows:

- Library 1 (Sample name in repository: S3): Feed control
- Library 2 (S4): Injection control
- Libraries 3 (S9) and 4 (S10): Feed WSSV infection
- Libraries 5 (S11) and 6 (S12): Injection WSSV infection.

Following library preparation, samples were sequenced by Novogene using Illumina NovaSeq X Plus, 150bp paired end sequencing (PE150) on the 25B flow cell. Libraries were spiked with 2% PhiX. Approximately 1.5 billion reads across six libraries were obtained, for an estimated 50,000 reads per cell.

The full sequencing dataset has been uploaded to the SRA database, under BioProject accession PRJNA1307069. The eight Parse sub-libraries can be found under BioSample accessions: SAMN50653283, SAMN50653284, SAMN50653285, SAMN50653286, SAMN50653287, SAMN50653288, SAMN50653289, SAMN50653290.

### 2.7 Data analysis

The data analysis was performed using a homemade pipeline (available: https://github.com/Roslin-Aquaculture/Parse-snRNAseq_LV-WSSV.git) that combined Parse Biosciences pipeline for Linux and the options from Seurat single cell analysis package (v4.4.0, Satija Lab, (23)) for R (v4.2.0, (24)). Below is an overview of the analysis steps undertaken. The full details, including individual sample thresholds for filtering can be found inside the GitHub repository.

#### 2.7.1 Data de-multiplexing

A genome file made from *Litopenaeus vannamei* (Genome assembly ASM378908v1, NCBI RefSeq assembly GCF_003789085.1) concatenated with the *White spot syndrome virus* (Genome assembly ASM397252v1, NCBI RefSeq assembly GCF_003972525.1) annotation was used in Parse split-pipe v1.2.1 (use_case “normal”; mode “comb”; chemistry “v2”; kit “WT_mega 96 wells”) to demultiplex sequence data into biological samples, align the reads against the genome, generate gene annotations, and estimate gene expression per cell, resulting in a unique molecular identifier (UMI) count matrix per sample.

#### 2.7.2 Cluster and Differential gene expression analysis using Seurat R package

The sample preprocessing, including quality controls (QC), was performed in Seurat v4.4.0 (SeuratObject v 4.1.4) as Seurat v4.3.0 or higher is required to open the ‘count_matrix.mtx’ file generated by Parse’s split-pipe. This included removing the cells with low UMI or feature numbers (detected genes), as well as those cells which contained more than 5% mitochondrial DNA genes (features). The UMI and feature thresholds vary with each sample and are specified inside the GitHub repository. This was followed by SCTransformation and cell cycle (st.flavor = “v2“; method = “glmGamPoi“; assay = “RNA”) data normalisation steps. The highly variable genes were identified (feature selection), the data was scaled accordingly, and a linear dimensional reduction was performed. Finally, the ‘dimensionality’ of the dataset was determined using Elbowplots (dims = 1:15), and the cells were clustered, generating the initial UMAP (Uniform Manifold Approximation and Projection) (resolution = 0.4).

After the initial UMAP, the DoubletFinder package was used to remove the doublets from the dataset. The dataset was integrated using integration anchors before downstream analysis. Individual UMAPS were generated per group (control or infected) and treatment (feed or injection). The cell distribution throughout the clusters was assessed based on group (control or infected) and treatment (feed or injection). From this point on Seurat versions v4.1.0 (Seurat Object 4.0.4) was used for all other analysis steps to accommodate for an already existing pipeline we developed for 10x Genomics data (available: https://github.com/Roslin-Aquaculture/10x-snRNAseq_LV-HP-atlas.git). Differential gene expression analysis (DEG) was performed in order to determine the gene markers for each cluster. At this point, a list containing the top 20 marker genes (by positive expression level) for each cluster was generated (assay = “SCT“; slot = “data“; only.pos = TRUE; min.pct = 0.25; return.thresh = 0.01; pseudocount.use = 0.001; logfc.threshold = 0.25; test.use = “LR“; latent.vars = “sample”). Each gene was analysed manually with a combination of NCBI (www.ncbi.nlm.nih.gov/gene/; (25)), UniProt (www.uniprot.org/; (26)) and The Human Protein Atlas (www.proteinatlas.org/; (27)). Following this, the UMAP clusters were annotated, and a series of plots (Heatmaps, Dotplots, Feature plots, Violin plots) were generated characterising each cluster and its relevant markers using Seurat.

Finally, expression differences between control and WSSV-infection groups by treatment were analysed using Seurat DEG analysis (Wilcoxon Rank Sum test using SCT assay; min.pct=0.1; return.thresh=0.01; pseudocount.use=0.001; logfc.threshold=0.25). The function of all the differentially expressed genes was assigned manually with a combination of NCBI (www.ncbi.nlm.nih.gov/gene/; (25)), UniProt (www.uniprot.org/; (26)) and The Human Protein Atlas (www.proteinatlas.org/; (27)). A Dotplot of the top 10 differentially expressed genes (by logFC value for both negative and positive values) by treatment was created per group and plotted DEGs were studied regardless of their inclusion in the Dotplots.

## 3. Results

### 3.1 WSSV infections and viral quantification

For the ‘WSSV injection’ group, the CT values were between 20.97 and 24.10, which indicates a viral load of between 3×10^6^ and 3×10^7^ WSSV particles per µl. For the ‘WSSV feed’ group, the CT values were between 22.41 and 29.32, indicating a viral load between 3×10^5^ and 1.5×10^7^ WSSV particles per µl (**Figure 4**).

**Figure 4.**
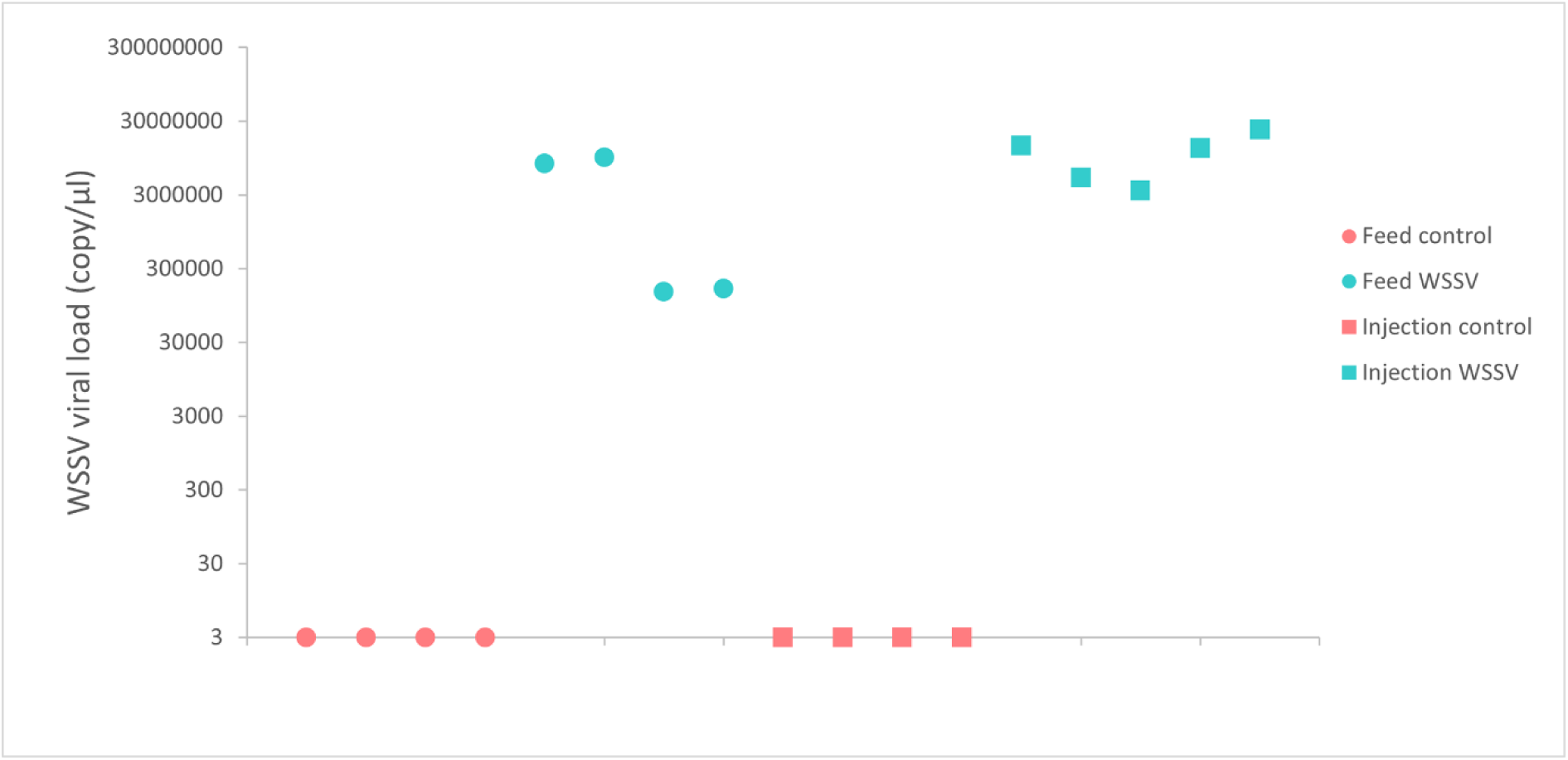
Average viral load of shrimp selected for snRNA sequencing analysis. The viral load was quantified using 50 mg of muscle tissue from the last abdominal segment. The viral load was inferred from the WSSV standard curve derived by qPCR (Ct=−3.7913x log (Concentration)+48.887).

### 3.2 Nuclei isolation

The lymphoid organ nuclei had minimal debris inside the final samples and had >95% singlet rate as counted using the LUNA-FX7™ Automated Cell Counter (**Figure 5**). The number of nuclei varied slightly between samples. The nuclei had minimal amount of degradation and were generally high quality,

**Figure 5.**
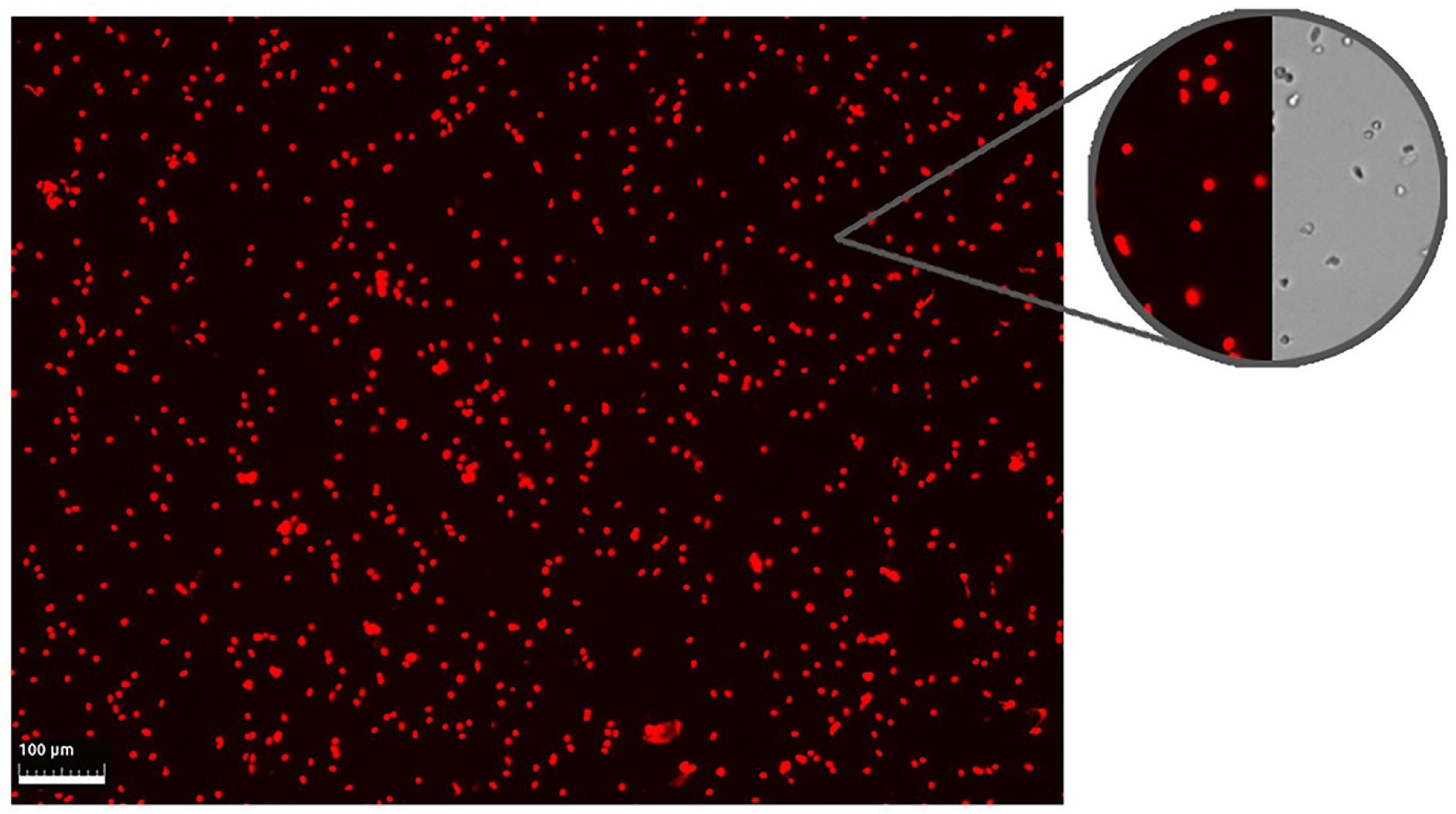
Example of isolated and fixated nuclei from lymphoid organ. The images are taken using the LUNA-FX7™ Automated Cell Counter. For the red field imaging, the nuclei were stained with Acridine Orange/Propidium Iodide dye. The top-right circles show a closer look at the nuclei in both red-field and bright-field

### 3.3 Cell atlas construction and histopathological data integration

All six snRNA-seq libraries passed filtration, with over 1.5 billion reads across all samples (0.89 sequencing saturation). The final number of cells obtained after the QC and filtering was 18,541 (**Figure 6 A**). The mean number of UMIs and genes was 1,173 UMIs and 654 feature per cell (**Figure 6 D, E**).

**Figure 6.**
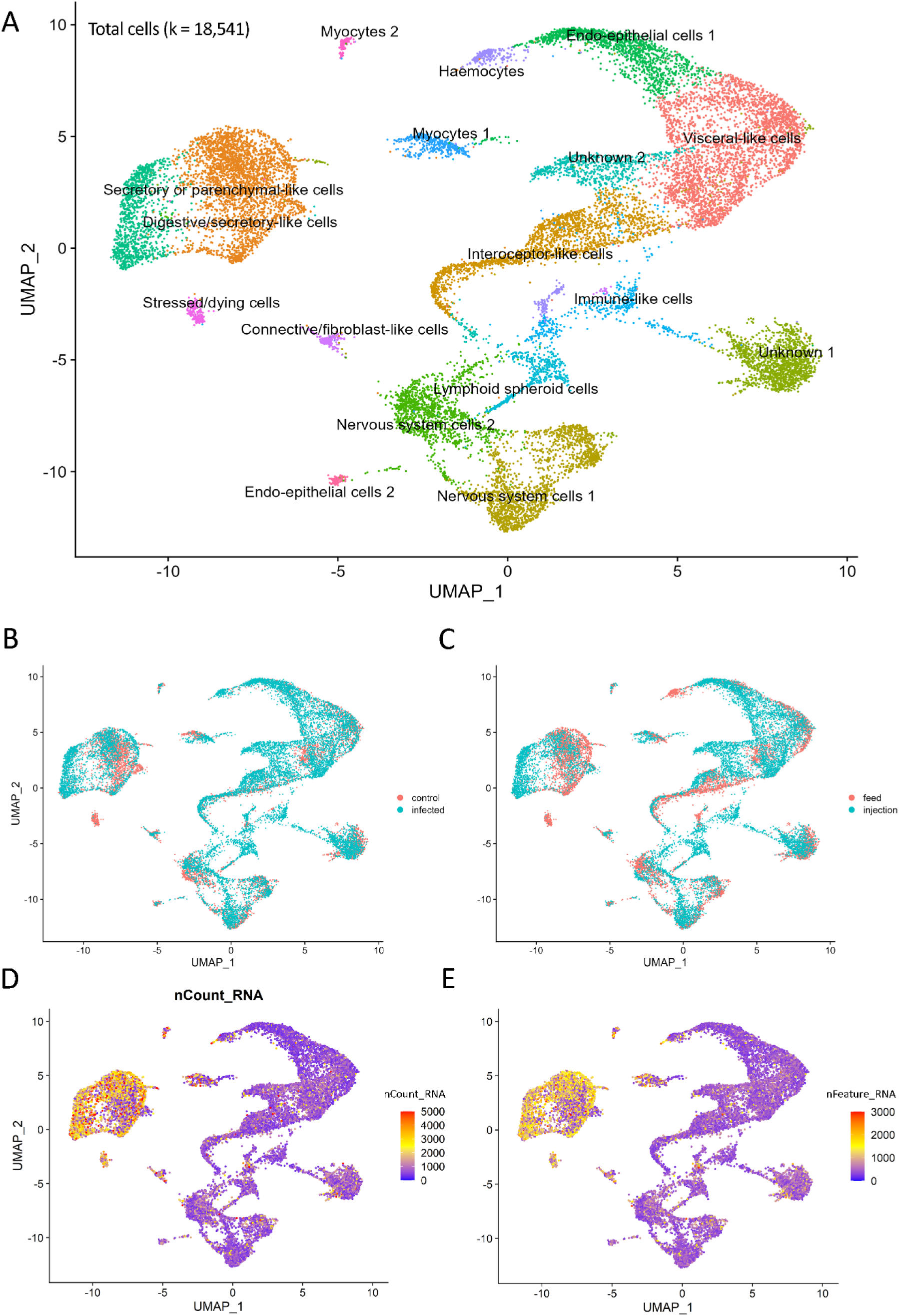
UMAPs. A) UMAP plot of cell clusters from shrimp lymphoid organ tissue. B) UMAP sample spread by treatment group (control and WSSV-infected). C) UMAP by infection method (feed vs injection). D) Total number of molecules (UMIs) detected within cells (nCount_RNA). E) Total number of genes detected in each cell (nFeature_RNA).

The nuclei number and distribution by sample group or treatment was also assessed. Overall, our experimental data contained 12,117 nuclei from WSSV-infected group and 6,424 from control (**Figure 6 B**), 11,284 nuclei from injection treatment and 7,257 from feed (**Figure 6 C**).

A total of 17 clusters were observed (**Figure 6 A**). The majority of clusters had distinct and recognisable transcriptomic profiles, with two clusters remaining undetermined due to the high number of uncharacterised marker genes and lack of cell type specificity of the annotated marker genes. The lymphoid organ cluster identification was partly based on the histopathological data from Rusaini and Owens (7).

#### 3.3.3 Lymphoid organ cell clusters

Distribution of the number of UMIs and features, as well as the percentage of UMIs that were mtDNA genes for each cluster can be seen in **Figure 7**. Marker genes differentiating clusters within the lymphoid organ can be seen in **Figure 8**. Here we aim to assign identity to each of the cell clusters based on expression of key and unique genes. As a reminder, the clusters were assigned based on the localisation and function of the top expressed genes (human or mouse homologues in most part) according to data from NCBI, UniProt and The Human Protein Atlas. Unless otherwise specified.

**Figure 7.**
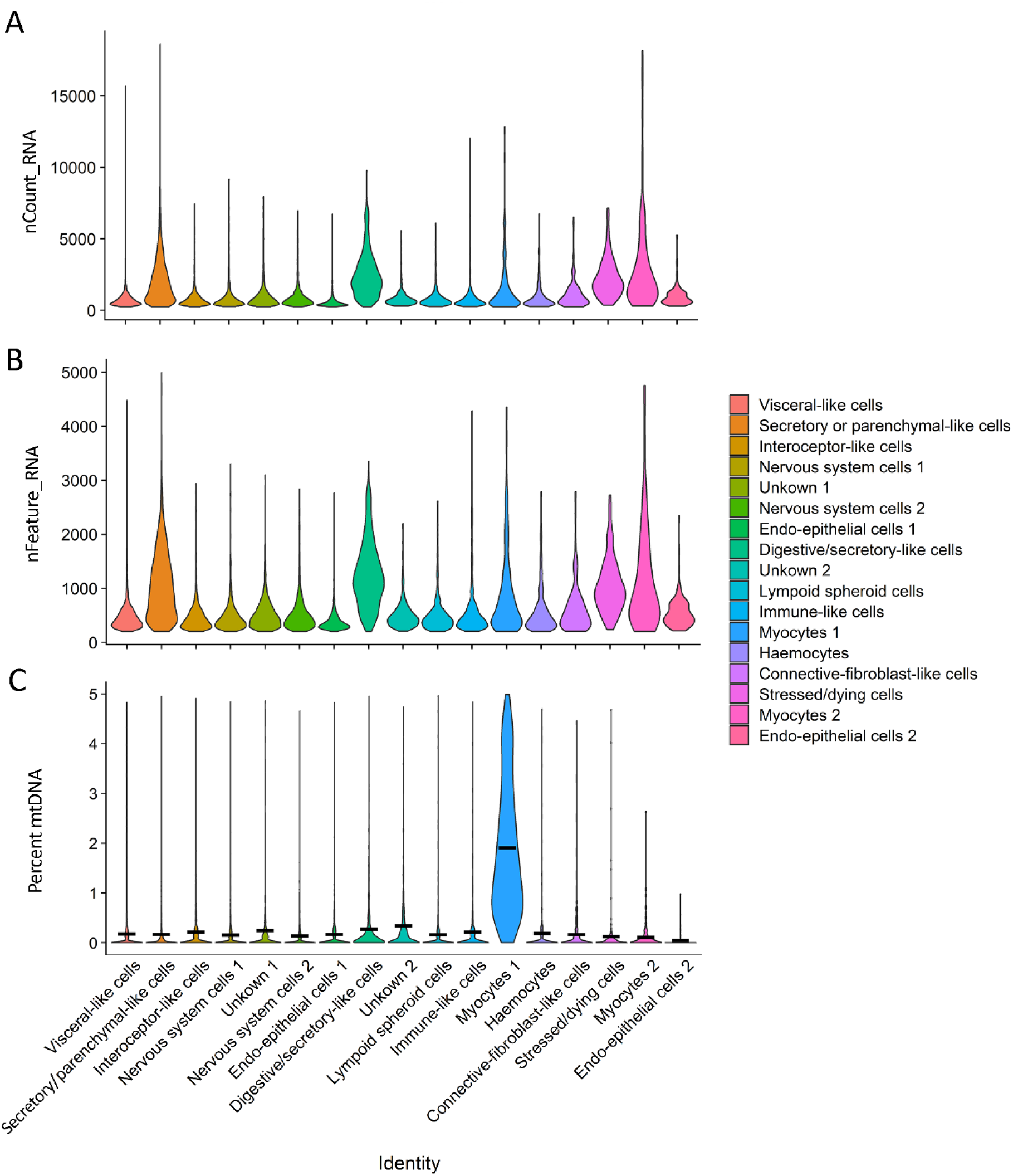
Data quality-violin plots by cluster. A) Average number of molecules (UMIs) detected within cells by cluster (nCount_RNA). B) Average number of genes detected in each cell by cluster (nFeature_RNA). C) Mitochondrial DNA percentage by cluster. The black bar represents average mtDNA percentage per cluster.

**Figure 8.**
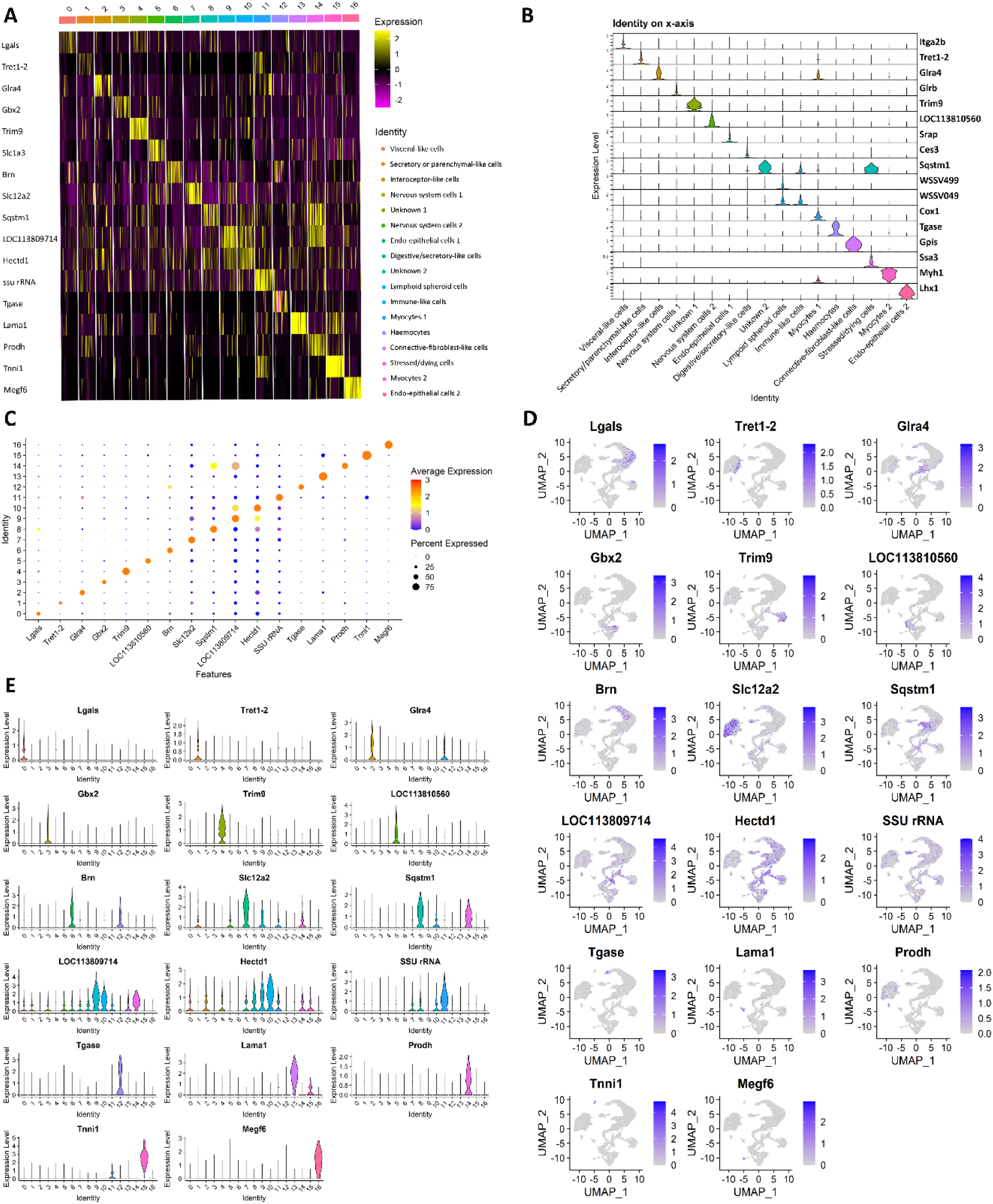
Cluster analysis. A) Heatmap of the top 8 marker genes in lymphoid organ. B) Violin plots of the significantly marker genes per cluster. C) Dot plot of significantly marker genes per cluster. Cluster numerical labels on the Y axis are as follows: 0 = Visceral-like cells, 1 = Secretory or parenchymal-like cells, 2 = Interoceptor-like cells, 3 = Nervous system cells 1, 4 = Unknown 1, 5 = Nervous system cells 2, 6 = Endo-epithelial cells 1, 7 = Digestive/secretory-like cells, 8 = Unknown 2, 9 = Lymphoid spheroid cells, 10 = Immune-like cells, 11 = Myocytes 1, 12 = Haemocytes, 13 = Connective-fibroblast-like cells, 14 = Stressed/dying cells, 15 = Myocytes 2, 16 =Endo-epithelial cells 2. D) UMAPs showing the expression level and location of the top cluster markers. E) Violin plots showing expression of top cluster markers by cluster.

##### Non-immune cell clusters

The clusters we identified include many of the cell types expected to be present within lymphoid tissues based on histopathological data present in the literature. We identified Cluster 0 as “Visceral-like cells” with low confidence due to a higher percentage of genes with low tissue specificity, and the presence of only a handful of genes (*Clca1, Ptprt, Pdgfrb*) indicative of the identity. It is worth nothing that there are several genes in this cluster with additional roles in hydrolysis and pro-inflammation, including *Lgals, Casp1, LyzPtprt, clca1* and *Fbh1*, but their localisation is very broad or non-specific. Nonetheless, it could indicate that this cluster may also be associated with immune-like cells such as macrophages instead. Cluster 1, “Secretory or parenchymal-like cells”, expresses genes involved in secretion, including transporters and solute carriers such as *Tret1-2, Slc5a12, Slc6a5, Naat1, Slco5a1, Slc5a8, Slc13a2*, and two homologues of *Slc22a1* and *Slco3a1*. Cluster 7 has a similar gene profile and was identified as “Digestive/Secretory-like cells” due to the presence of *Slc12a2, Clca1, Naat1, Ces3* and *Ces2a*, often seen expressed in digestive glands due to their roles in transport (ion, calcium, sodium, potassium, amino-acid transporters) and hydrolysis. Cluster 2 expresses a mix of receptor-coding genes (*Glra4, Scarb1, Hr96*), generally enriched in internal-organ cells, as well as genes coding for proteins involved in neurogenesis (*Dar1*) and gap-junction structure (*Inx2*). As such, the cells belonging to this cluster are identified as potential interceptors (sensory receptor for internal organs). Cluster 3 and Cluster 5 are both identified as nervous system cell types with high confidence due to the expression of CNS and PNS-enriched genes. Cluster 3 expressed *Gbx2, Hth,* two homologues of *Cadn*, and *Ap,* which have functions in DNA binding and/or cell adhesion inside the brain, bone marrow and peripheral neurons and *Glrb* which codes for glycine receptors (in ligand-gated chloride channels). Cluster 5 expresses *Slc1a3* (transporter enriched in astrocytes)*, Nrxn1* (neuronal cell surface protein with role in cell adhesion)*, Grik2* (receptor for the excitatory neurotransmitter L-glutamate) and *Ten-M* (cell adhesion role in neural development) among its top 20 gene markers. Although we cannot assign a definitive identity to Cluster 4, multiple embryonic-enriched genes such as *Rst*, *If,* and two copies of *Trim9*, indicate an undifferentiated cluster of cells. The “Endo-epithelial cells 1” cluster (Cluster 6) has gene markers enriched in both epithelial tissue (*Brn, Notch1, Mys*) as well as a variety of genes that are typically enriched in internal organs lining or tubes in vertebrates. An additional smaller cluster of “Endo-epithelial cells 2” (Cluster 17) was also identified, although we predict that this cluster might contain undifferentiated cells, possibly similar to pluripotent stem cells. This is due to the presence of *Lhx5*, two homologues of *Lhx1*, two LIM/homeobox protein-coding genes that are enriched in the developing vertebrate brain and kidney, as well as a mix of genes that are generally expressed in endothelial or epithelial-like cells from a multitude of organs. Cluster 8 remains uncharacterised due to half of the top 20 markers belonging to uncharacterised loci, as well as the low tissue expression specificity of the characterised markers. Clusters 11 and 15 are both myocytes. Both clusters express multiple copies of *Ttn, Tnni1* and *Myh1*. In addition, Cluster 11 expresses *Mt-Nd3* and Cluster 15 expresses *Tnnc1* as a marker. We note that Cluster 11 has an unusually high proportion of mitochondrial DNA comparing to the other clusters (including *Cox1* gene as a marker) (Figure 7C), in addition to three RNA-specific genes coding for large and small RNA subunits (one SSU and two LSU rRNA genes), which could be indicative of a poorer quality cluster. Cluster 13 has gene markers for different cell types and was difficult to assign to a single identity. Nevertheless, there were several genes (*Lama4, Angptl2, Reln*) that indicated it may be formed of connective tissue or fibroblast-like cells. Lastly, Cluster 14 is the only cluster to be made up of cells entirely from the control group samples. This cluster expresses a variety of heat-shock or heat-shock-binding protein-encoding genes: *Hsc70, Hsc70-4, Dnajb4, Ssa3, Hsp20* and *Hspbp1*. These types of proteins maintain cellular proteostasis and protect cells from stresses (28). As such, we concluded that this cluster is likely stressed or dying cells.

##### Immune cell clusters

Clusters 9 and 10 were made almost entirely out of cells from the WSSV infection samples, potentially indicating an influx of these particular cell types into the lymphoid organ at the time of infection. Additionally, these clusters were the only two to contain WSSV genes as markers. Cluster 9 is likely to represent lymphoid organ-specific spheroid cells. Nine out of the top 20 markers were WSSV genes. Alongside them, multiple *L. vannamei* genes were expressed, including *Abcc1*, a membrane pump which mediates the export of foreign material from the cytoplasm, *Cpne8*, which regulate molecular events at the interface of the cell membrane and cytoplasm, as well as two cancer and tumour related genes: *Brat* and *Unc5c*. Similarly, Cluster 10 had two WSSV genes as markers. In addition to that, it expressed genes generally associated with immune cells (plasma and NK cells) such as *Npc1* and *LpiN3*, which suggest the cluster is of “Immune-like cells”. Furthermore, we also observed the presence of several genes involved in early cell development and embryogenesis, meaning that this cluster could potentially be young spheroid or young immune (haemocyte) cells. Finally, Cluster 12 has been identified as haemocytes due to the expression of the hemocytin-like gene, *proclotting enzyme-like* gene and two *Tgase* homologues.

### 3.4 Differential gene expression (DEGs) analysis

Across the whole dataset (**Figure 9**, **Figure 10**), we found 85 upregulated and 70 downregulated genes in the WSSV-infected group. From these, 13 upregulated genes and 19 downregulated genes had an average log fold change of over 5.0, and there were three genes with values over 9.0. Several differentially expressed genes are likely to play a role in viral immunity or apoptosis. *Slc2a1*, a viral receptor in humans was upregulated in the “Myocytes 2” cluster (avg_log2FC = 5.71) but also downregulated in “Endo-epithelial cells 1” cluster (avg_log2FC = −4.39). Two different copies of *Abcc5* gene were downregulated in the “Myocytes 2” cluster (avg_log2FC = −2.87 and avg_log2FC −4.50). *Ivns1abp*, an antiviral-related gene, was downregulated in the “Secretory/parenchymal-like cells” cluster (avg_log2FC = −4.71). Two genes involved in host-virus interactions were also found among the differentially expressed genes. *Xpo1* (avg_log2FC = 3.71) was upregulated in the “Connective tissue-fibroblast-like” cluster while *Cebpa* (avg_log2FC = −2.65) was downregulated in the” Endo-epithelial cells 1” cluster. Apoptosis-related genes such as *Slk* (avg_log2FC = 4.82) and *Plxna3* (avg_log2FC = 4.15) were upregulated in the “Connective -fibroblast-like” and “Visceral-like cells” clusters respectively, while others such as *Ptpn4* (avg_log2FC = −2.05), *Ntrk2* (avg_log2FC = −6.72) and *Comp* (avg_log2FC = −6.88) were downregulated in the “Nervous system cells 2”, “Lymphoid spheroid cells” and “Endo-epithelial cells 1” cluster respectively. *Hr3*, a putative receptor, was the only differentially expressed gene present in both tissues. This gene was downregulated in the “Nervous system cells 2” cluster (avg_log2FC = −3.92).

**Figure 9.**
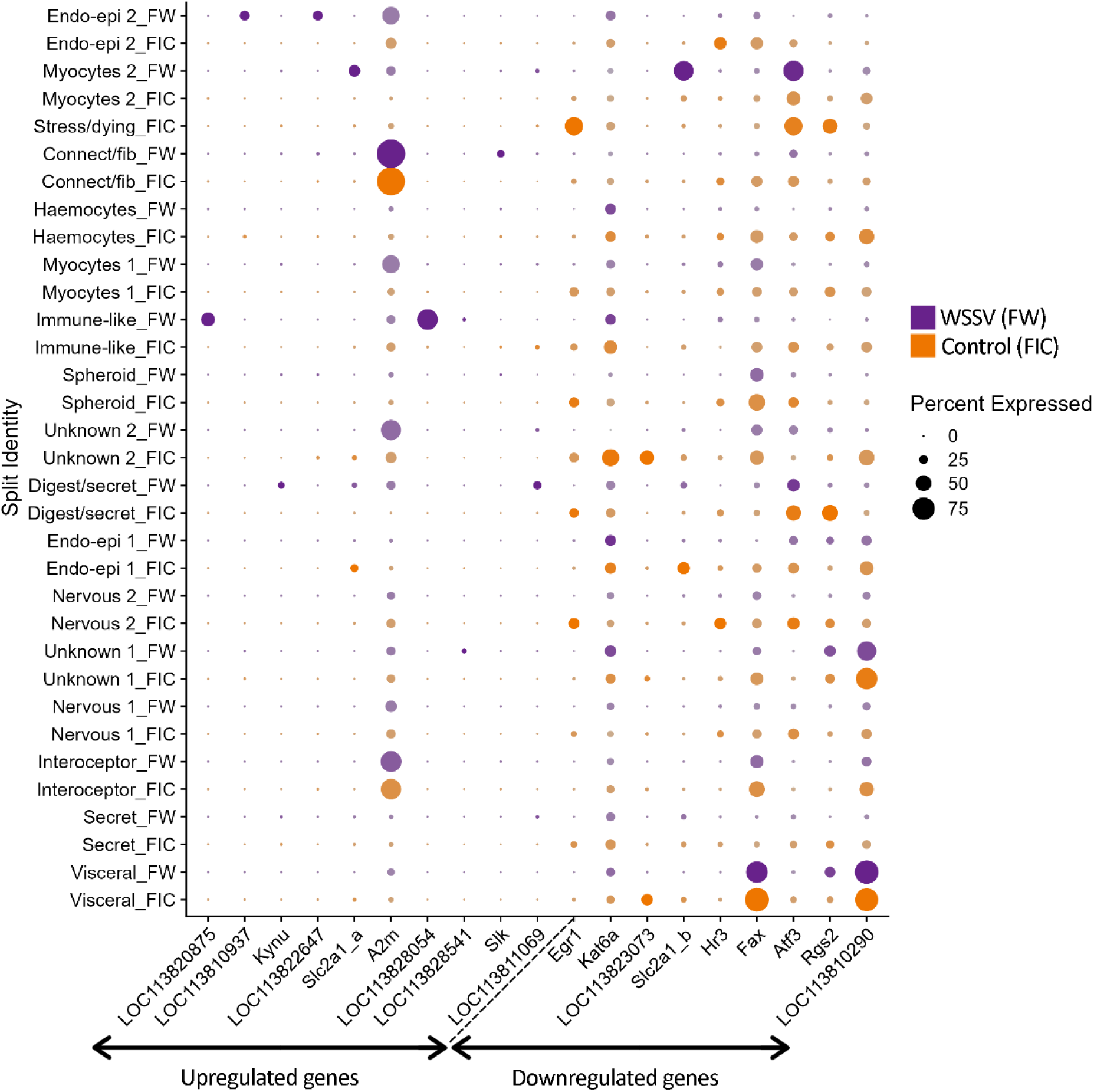
Differential gene expression dotplot in the ‘Feed’ group. The plots show the differentially expressed genes between the control and WSSV-infection samples in the ‘Feed’ infection group. The top 10 upregulated and top 10 downregulated genes per infection method were chosen for these plots. The size of the dots coincides with the gene expression percentage while the different colours represent the control or infected groups. The colour intensity coincides with the difference in expression levels, with the most intense colour representing up to 9.89 times expression level difference.

**Figure 10.**
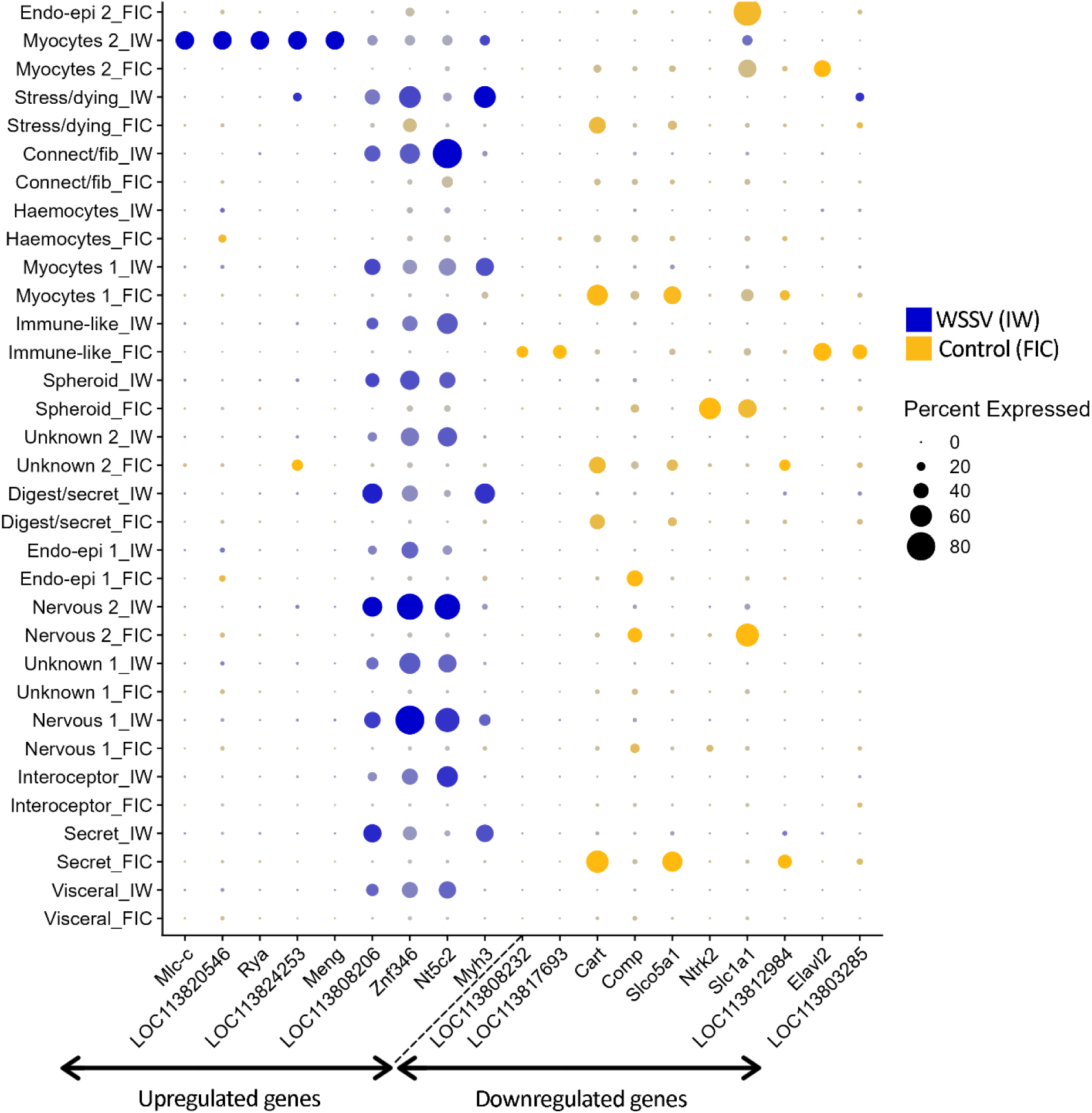
Differential gene expression dotplot in the ‘Injection’ group. The plots show the differentially expressed genes between the control and WSSV-infection samples in the ‘Injection’ infection group. The top 10 upregulated and top 10 downregulated genes per infection method were chosen for these plots. The size of the dots coincides with the gene expression percentage while the different colours represent the control or infected groups. The colour intensity coincides with the difference in expression levels, with the most intense colour representing up to 9.89 times expression level difference.

## 4. Discussion

In this study, we conducted a white spot syndrome virus challenge on Pacific whiteleg shrimp in order to understand the molecular mechanisms underlying both the host response to WSSV and how the virus modulates gene expression in host cells. All WSSV-challenged animals exhibited clinical symptoms associated with white spot disease. In the case of acute infections (“Injection” group), the shrimp only exhibited lethargy, reduction in feeding and slight red discoloration on the body. The shrimp belonging to the “Feed” group also developed white spots on the exoskeleton, the “hallmark” of WSD, due to the more chronic nature of this infection method, as seen in other penaeid shrimp such as *P. indicus* (29).

### 4.1 Cell types found in the lymphoid organ

The lymphoid organ is a paired, lobed, immune organ and a crucial component of the immune system, exhibiting significant differential expression of genes involved in viral receptor activity, immune modulation, cytokine production, pathogen-host interactions, cellular stress responses and apoptosis (Rusaini and Owens, 2010);(11). It is involved in cellular and humoral immune responses and acts as the primary site of viral degradation via its spheroid cells (10); (11). Located ventro-anterior to the hepatopancreas, it is structurally characterized by lymphoid tubules and hemal sinuses, where it functions primarily as a haemolymph filter. The two lobes are surrounded by connective tissue capsules and muscle, and are connected to the heart by the subgastric artery. The tubules are surrounded by connective tissue fibre, have a central haemal lumen, stromal matrix cells and interstitial sinuses. The lumen is lined with flattened endothelial cells and often blocked with haemocytes (7).

We have successfully developed a new protocol for nuclei isolation from this important organ which will crucially inform future snRNA-seq studies on the immunological role of this organ. We have also generated a detailed cell atlas of the lymphoid organ, identifying multiple cell types with distinct roles in both function and viral immunity.

Our samples contained various clusters with structural supportive or motility role, including epithelial cells (Clusters 6 and 16), muscle cells (Clusters 11 and 15) and connective tissue cells (Cluster 13). Three distinct clusters with roles in secretion were also identified (Cluster 0, 1 and 7). In vertebrates, lymphoid secretory cells are known to play important roles in immune function by facilitating communication, pathogen defences, and tissue organization (30). While lymphoid tissues (in vertebrates) are not primarily sensory organs, they do contain nervous elements (such as autonomic nerve fibres and sensory nerve endings) that can influence immune function and that provide a major pathway through which the brain can alter immune reactivity (31). We identified three different clusters that potentially fulfil these roles (Clusters 2,3 and 5).

Lymphoid organ plays a key role in pathogen defence by facilitating the recognition and elimination of viral and bacterial invaders and has a significant role during pathogenic infections (7); (11). The spheroid cells (Cluster 9) are formed in the haemal sinuses and appear to have a more basophilic cytoplasm and lack of a central lumen as compared to normal lymphoid tubules (7). The spheroid cells phagocytize foreign material, particularly viruses (10). These cells form during infection, encapsulate the pathogen and then degenerate. They are known to exhibit anaplasia (characteristic of malignant tumours) and to have an increased cytoplasm to nuclear ratio (7).

### 4.2 Infection- and immunity-related DEGs in lymphoid organ (**Table 3**)

**Note for Editors: Please insert Table 3 here (found at the end of the document due to formatting)**

**Table 3.**
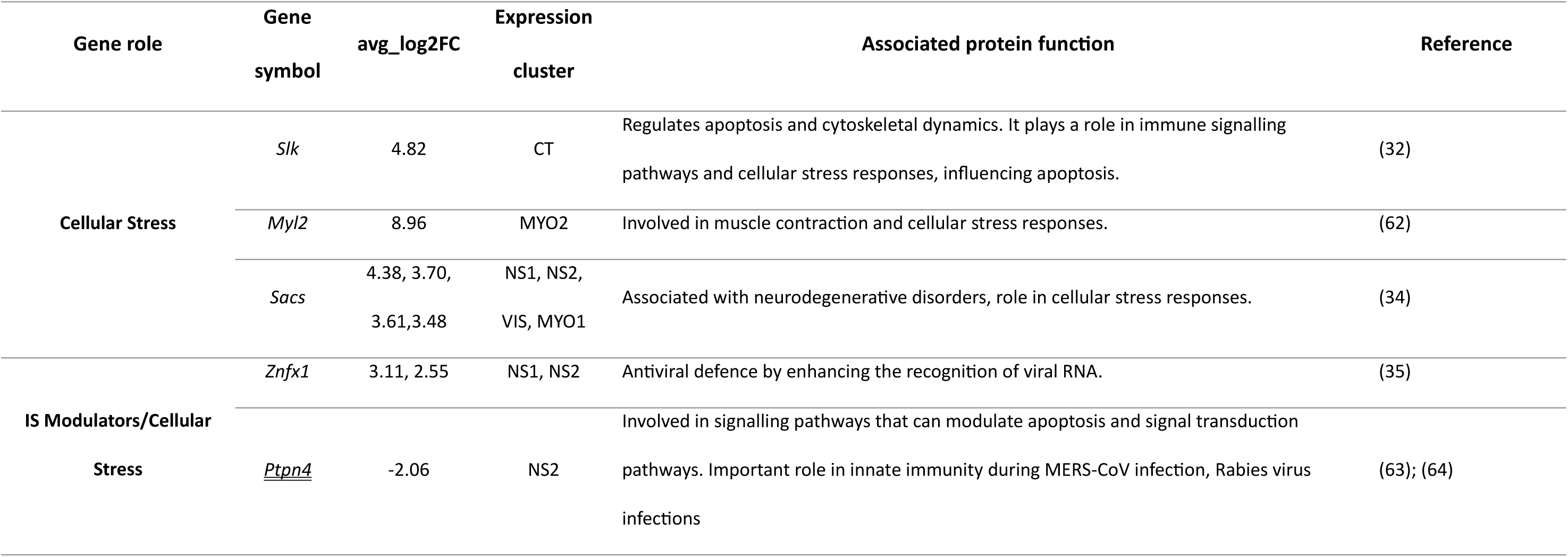
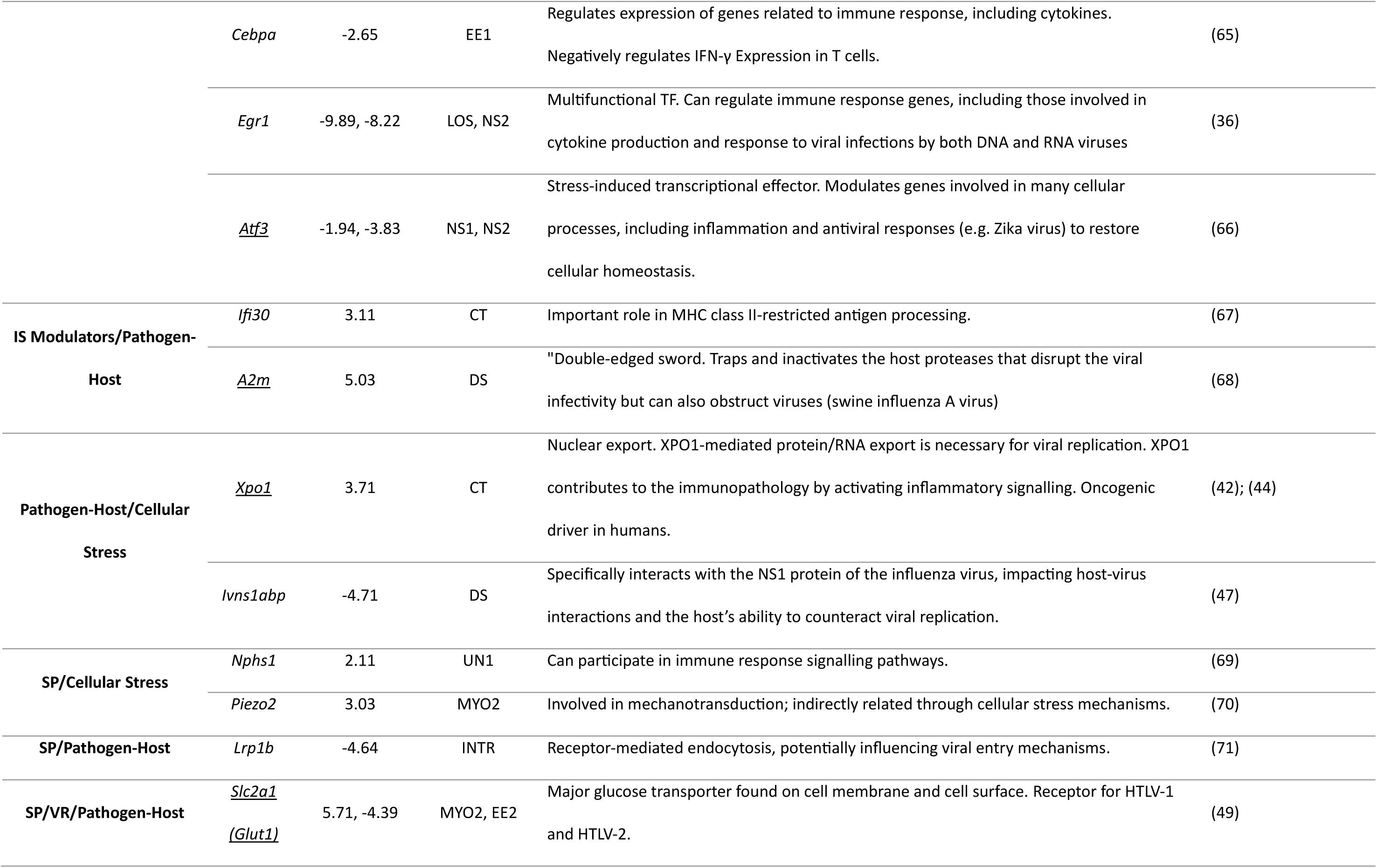
Immune- and infection-related genes differentially expressed in lymphoid organ. TF=transcription factor, VR=Viral receptor, IS=Immune system, INTR=Interoceptor-like cells, CT= connective fibroblasts, MYO=Myocytes, DS=Digestive/secretory-like cells, LOS=Lymphoid spheroid cells, VIS=Visceral-like cells, UN=Unknown, EE=Endo-epithelial cells, NS=Nervous system cells. The gene relative expression level between control and WSSV-infected samples is shown as avg_log2FC, where positive numbers represent upregulated genes in infected samples, and negative numbers represent downregulated genes. All the functions stated have been primarily studied in mammalians, unless stated otherwise. All underlined genes have also been studied in invertebrates. The gene underlined with a double line was differentially expressed in both tissues.

Two genes associated with cellular stress response, *Slk* and *Sacs*, were found to be differentially expressed in our data. *Slk* drives protein homodimerization activity and protein serine/threonine kinase activity and plays a role in immune signalling pathways and cellular stress responses (32). The gene also plays a role in apoptotic signalling pathways, which are crucial for the elimination of infected or damaged cells. The kinase activity of SLK is involved in the activation of stress-activated protein kinase (SAPK) pathways, such as the JNK and p38 MAPK pathways, which are involved in immune responses and inflammation (33). *Sacs*, though not directly related to viral resistance, may influence cellular stress responses and overall immune function. *Sacs* has chaperon activities, controlling the microtubule balance or cell migration and has been associated with neurodegenerative diseases (34). SACS contains domains similar to heat shock proteins and J-domain proteins, which are known to participate in stress responses. This function may influence how cells manage protein misfolding during viral infections, which can lead to stress and aggregation of viral proteins.

*Znfx1, Egr1, Atf3* and *A2m* code for immune system modulators that also have a role in cellular stress. ZNFX1 is a zinc finger protein involved in antigen processing. It plays multiple roles in immune response modulation. It is an interferon-stimulated double-stranded RNA sensor that recognises viral RNA and restricts the replication of RNA viruses in mice (35), highlighting its potential in enhancing immune recognition against WSSV. EGR1 is involved in cytokine production during viral infections of both DNA and RNA viruses. In mice, several cytokines, including IFNα, INFβ, IFNλ, and TNFα, are capable of increasing *Egr1* transcription (36). In our study, *Egr1* was highly downregulated in two different cell types, potentially indicating host-cell manipulation by WSSV to create a suitable environment for replication. An additional immune system modulator is ATF3, which is a stress-induced transcriptional effector shown to be induced by viral infections such as the Human cytomegalovirus (HCMV) (37) and Zika virus (ZIKV) in human cells. Additionally, *Atf3* is induced in cells infected with Herpes Simplex Virus (HSV) and helps maintain the virus in a latent (resting) state. It can influence the cellular environment to either promote or inhibit viral replication (38). In invertebrates, this gene has been characterized in silkworms (*Bombyx mori*) and studied more extensively in *Drosophila*. In the latter species, *Atf3* plays a role in cellular stress responses and immune signalling, safeguarding metabolic and immune system homeostasis (39). Lastly among immune modulators we highlight A2M which is involved in pathogen-host interactions. A2M, a broad-spectrum protease inhibitor, may play a role in neutralizing WSSV proteases, thereby inhibiting viral spread within the host (40).

XPO1 is crucial for the nuclear export of viral ribonucleoproteins in viruses like HIV (Yarbrough Boons et al., 2015), Influenza (42), and Ebola (43). In humans, XPO1 also contributes to immunopathology by activating inflammatory signalling and is an oncogenic driver (44). In *Drosophila, Xpo1* has been studied for its role in nuclear protein and RNA transport which influences developmental and stress response pathways (45). In crustaceans, modulation of *Xpo1* could potentially disrupt WSSV replication, much like it does during HIV infection, where the inhibition of this gene suppresses the replication of HIV and induces apoptosis in infected cells.

*Ivns1abp* is involved in host-pathogen interactions, particularly with influenza viruses. In humans, *Ivns1abp* encodes a protein that interacts with the non-structural protein 1 (NS1) of influenza A viruses (IAV) (46). The NS1 protein of influenza viruses plays crucial roles in evading host immune responses and facilitating viral replication within host cells (47). In our case, this gene was downregulated in infected samples, potentially as an immunity avoidance mechanism mounted by WSSV.

One final gene that is linked to viral infections is *Slc2a1*, also known as *Glut1*, which codes for a cell membrane glucose transporter implicated in viral entry and replication processes in various viruses such as HIV (48), as well as acting as a cell surface receptor for viral entry in Human T-lymphotropic virus 1 (HTLV-1) and HTLV-2 infections (49). This gene has also been studied in various invertebrate species, particularly in the context of metabolic and stress responses. It is involved in nutrient transport processes that are crucial for energy balance in invertebrates, such as *D. melanogaster,* and it has been shown to be an important gene in the outcome of Huntington’s disease where an increase in gene expression leads to ameliorated disease phenotype (50). In crustaceans, *Glut1* expression is important for hemocytin adaptation to hypoxic conditions in the Oriental River prawn (*Macrobrachium nipponense*) (51). In Pacific whiteleg shrimp, Glut1 was shown to be a conserved protein whose expression changes with dietary carbohydrate level and source, as well as environmental stressors such as salinity, temperature changes and hypoxia (52); (53). Although its direct role in crustacean viral infections remains to be determined, the differential control of the *Slc2a1* among the different lymphoid organ cell clusters suggests its potential involvement in both host defence and WSSV entry and replication, making it an important gene to study in the context of infections and immunity.

### 4.3 WSSV infection mechanism and related genes

White spot syndrome (WSS) is one of the most damaging viral diseases in crustacean aquaculture, displaying aggressive pathogenicity that can result in up to 100% mortality 3-10 days post-infection (54). WSSV’s life cycle can be divided into three steps: entry into the host cell, uncoating of the genome followed by replication, and, particle assembly and release (55); (56). The first step of WSSV’s cycle is the entry into host cells. This is done by binding to specific receptor molecules on the surface of target cell, which leads to induction of Clathrin-mediated endocytosis. Both LRP1B and SLC2A1 could potentially serve this purpose. Although LRP1B is not studied as a specific receptor, members of the LDL receptor family, including LRP1, are known to be exploited by viruses like Hepatitis C virus (HCV) for cell entry (57);(58). GLUT1 (*slc2a1*) has been identified as a receptor for HTLV-1 and HTLV-2, facilitating the virus’s entry into host cells (49).

Once in the cell, host transcription factors, such as *Ie1,* bind the WSSV genome and initiate viral gene expression (59). WSSV establishes a IE1/JNK/c-Jun positive feedback loop to drive its replication (59). The SLK kinase is one of the molecules involved in the activation JNK pathway (33). Other several genes can come into play for the replication step such as *Gsk3b, Cebpa,* and *Atf3*. GSK3β is known to be manipulated by Human cytomegalovirus (HCMV) to create a host cellular environment geared towards accelerated protein translation support viral replication (60). CEBPA is involved in regulating immune responses and has been shown to interact with viral proteins from Hepatitis B virus (HBV), affecting viral replication via transcriptional regulation of gene expression (61). The modulation of *Atf3* by the virus could promote or inhibit viral replication by controlling the switch between the latent and active phases of the virus, as in the case of HSV (38). Lastly, after the viral genome replication, WSSV will use the cell machinery to assemble new viral particles and release them. XPO1 is used by many viruses, including SARS-CoV-2, MERS-CoV, HIV, Influenza, and Ebola, to export viral RNA and proteins from the nucleus to the cytoplasm, which is essential for viral replication and assembly (42). EGR1 is known to enhance viral gene expression, viral replication, and the release of infectious particles (36).

## 5. Conclusion

This study employed single nuclei RNA sequencing transcriptomic analysis of WSSV-infected adult Pacific whiteleg shrimp to uncover genes involved in viral immunity and infection. We generated novel WSSV disease challenge protocols for penaeid shrimp WSSV as well as a novel nuclei isolation protocol for the shrimp lymphoid organ, a critically-important immune organ. The sequencing data we generated was used to create a cell atlas for the lymphoid organ. The differential expression of genes involved in viral entry, immune modulation, pathogen-host interactions, and cellular stress responses underscores the complexity of the shrimp’s immune response to WSSV.

Our research highlighted a selection of WSSV infection-related genes. Future studies should focus on validating these genes through functional assays and exploring their roles in the broader context of crustacean immunity. This could potentially create a future path towards the creation of WSSV-resistant shrimp stocks via CRISPR gene editing and selective breeding, thereby enhancing aquaculture sustainability and productivity.

## List of abbreviations

Cas9: CRISPR-Associated Protein 9
CRISPR: Clustered Regularly Interspaced Short Palindromic Repeats
DEG: Differentially Expressed Genes
DNA: Deoxyribonucleic Acid
HCMV: Human Cytomegalovirus
HCV: Hepatitis C virus
HIV: Human Immunodeficiency Virus
HTLV: Human T-Lymphotropic Virus
IAV: Influenza A Viruses
IFN: Interferon
JNK: c-Jun N-terminal kinases
LIM (in LIM/homeobox): Lin-11, Islet-1, and Mec-3 - (the three original members of the LIM protein family)
LSU rRNA: Large Ribosomal RNA Subunit
MAPK: Mitogen-Activated Protein Kinases
MERS-CoV: East Respiratory Syndrome Coronavirus
NK: Natural Killer Cells
QPCR: Quantitative Polymerase Chain Reaction
RNA: Ribonucleic Acid
SAPK: Stress-Activated Protein Kinase
SARS-CoV-2: Severe-Acute-Respiratory-Syndrome-Related Coronavirus 2
snRNA-seq: Single Nuclei RNA Sequencing
SPF: Specific Pathogen-Free
SSU rRNA: Small Ribosomal RNA Subunit
TNF: Tumor Necrosis Factor
UMI: Unique Molecular Identifier
ZIKV: Zika Virus

## Availability of data and materials

The datasets analyzed during the current study are available from NCBI BioProject database (Project accession PRJNA1307069) under BioSample accessions: SAMN50653283, SAMN50653284, SAMN50653285, SAMN50653286, SAMN50653287, SAMN50653288, SAMN50653289, SAMN50653290.

## Ethics approval and consent to participate

This study involved the use of Pacific whiteleg shrimp (*Litopenaeus vannamei*) which are not currently regulated under A(SP)A 1986 but have recently become recognized under the Animal Welfare (Sentience) Act 2022. This study has been reviewed and approved by the Roslin Institute Animal Welfare and Ethical Review Body (AWERB).

## Acknowledgements

We would like to thank Andrew Whiston from RasTech for helping in the construction of our RAS system and for providing advice on shrimp husbandry and RAS system setup.

## References

1. FAO. The State of World Fisheries and Aquaculture 2024. FAO; 2024.

2. Flegel TW. A future vision for disease control in shrimp aquaculture. J World Aquac Soc. 2019 Apr 7;50(2):249–66.

3. Sánchez-Paz A. White spot syndrome virus: an overview on an emergent concern. Vet Res. 2010 Nov 26;41(6):43.

4. Huang Z, Aweya JJ, Zhu C, Tran NT, Hong Y, Li S, et al. Modulation of Crustacean Innate Immune Response by Amino Acids and Their Metabolites: Inferences From Other Species. Front Immunol. 2020 Nov 5;11.

5. Tassanakajon A, Somboonwiwat K, Supungul P, Tang S. Discovery of immune molecules and their crucial functions in shrimp immunity. Fish Shellfish Immunol. 2013 Apr;34(4):954–67.

6. Koiwai K, Kondo H, Hirono I. scRNA-seq analysis of hemocytes of penaeid shrimp under virus infection. Cold Spring Harbor Laboratory. 2023 Jan 4;

7. Rusaini, Owens L. Insight into the lymphoid organ of penaeid prawns: A review. Fish Shellfish Immunol. 2010 Sep;29(3):367–77.

8. Cao J, Wu L, Jin M, Li T, Hui K, Ren Q. Transcriptome profiling of the Macrobrachium rosenbergii lymphoid organ under the white spot syndrome virus challenge. Fish Shellfish Immunol. 2017 Aug;67:27–39.

9. Du ZQ. Comparative transcriptome analysis reveals three potential antiviral signaling pathways in lymph organ tissue of the red swamp crayfish, Procambarus clarkii. Genetics and Molecular Research. 2016;15(4).

10. Anggraeni M, Owens L. The haemocytic origin of lymphoid organ spheroid cells in the penaeid prawn Penaeus monodon. Dis Aquat Organ. 2000;40:85–92.

11. Wang F, Li S, Li F. Different Immune Responses of the Lymphoid Organ in Shrimp at Early Challenge Stage of Vibrio parahaemolyticus and WSSV. Animals. 2021 Jul 21;11(8):2160.

12. Khan YS, Farhana A. Histology, cell. StatPearls - NCBI Bookshelf. 2023;

13. Cui C, Tang X, Xing J, Sheng X, Chi H, Zhan W. Single-cell RNA-seq revealed heterogeneous responses and functional differentiation of hemocytes against white spot syndrome virus infection in *Litopenaeus vannamei*. J Virol. 2024 Mar 19;98(3).

14. Daniels RR, Taylor RS, Robledo D, Macqueen DJ. Single cell genomics as a transformative approach for aquaculture research and innovation. Rev Aquac. 2023 Sep 7;15(4):1618–37.

15. Salisbury SJ, Daniels RR, Monaghan SJ, Bron JE, Villamayor PR, Gervais O, et al. Keratinocytes drive the epithelial hyperplasia key to sea lice resistance in coho salmon. BMC Biol. 2024 Jul 29;22(1):160.

16. Cui C, Tang X, Xing J, Sheng X, Chi H, Zhan W. Single-cell RNA-seq uncovered hemocyte functional subtypes and their differentiational characteristics and connectivity with morphological subpopulations in Litopenaeus vannamei. Front Immunol. 2022 Sep 13;13.

17. Yang P, Chen Y, Huang Z, Xia H, Cheng L, Wu H, et al. Single-cell RNA sequencing analysis of shrimp immune cells identifies macrophage-like phagocytes. Elife. 2022 Oct 6;11.

18. Li Y, Zhou F, Yang Q, Jiang S, Huang J, Yang L, et al. Single-Cell Sequencing Reveals Types of Hepatopancreatic Cells and Haemocytes in Black Tiger Shrimp (Penaeus monodon) and Their Molecular Responses to Ammonia Stress. Front Immunol. 2022 May 4;13.

19. Zhu W, Yang C, Chen X, Liu Q, Li Q, Peng M, et al. Single-Cell Ribonucleic Acid Sequencing Clarifies Cold Tolerance Mechanisms in the Pacific White Shrimp (Litopenaeus Vannamei). Front Genet. 2022 Jan 12;12.

20. Koiwai K, Koyama T, Tsuda S, Toyoda A, Kikuchi K, Suzuki H, et al. Single-cell RNA-seq analysis reveals penaeid shrimp hemocyte subpopulations and cell differentiation process. Elife. 2021 Jun 16;10.

21. WOAH. Manual of diagnostic tests for aquatic animals. World Organisation for Animal Health; 2016.

22. Ruiz Daniels R, Taylor RS, Dobie R, Salisbury S, Furniss JJ, Clark E, et al. A versatile nuclei extraction protocol for single nucleus sequencing in non-model species–Optimization in various Atlantic salmon tissues. PLoS One. 2023 Sep 7;18(9):e0285020.

23. Hao Y, Hao S, Andersen-Nissen E, Mauck WM, Zheng S, Butler A, et al. Integrated analysis of multimodal single-cell data. Cell. 2021 Jun;184(13):3573–3587.e29.

24. R Core Team. R Foundation for Statistical Computing. Vienna, Austria. URL https://www.r-project.org/. 2021. R: A language and environment for statistical computing..

25. Sayers EW, Beck J, Bolton EE, Brister JR, Chan J, Connor R, et al. Database resources of the National Center for Biotechnology Information in 2025. Nucleic Acids Res. 2025 Jan 6;53(D1):D20–9.

26. Bateman A, Martin MJ, Orchard S, Magrane M, Adesina A, Ahmad S, et al. UniProt: the Universal Protein Knowledgebase in 2025. Nucleic Acids Res. 2025 Jan 6;53(D1):D609–17.

27. Uhlén M, Fagerberg L, Hallström BM, Lindskog C, Oksvold P, Mardinoglu A, et al. Tissue-based map of the human proteome. Science (1979). 2015 Jan 23;347(6220).

28. Hu C, Yang J, Qi Z, Wu H, Wang B, Zou F, et al. Heat shock proteins: Biological functions, pathological roles, and therapeutic opportunities. MedComm (Beijing). 2022 Sep 2;3(3).

29. Pradeep B, Rai P, Mohan SA, Shekhar MS, Karunasagar I. Biology, Host Range, Pathogenesis and Diagnosis of White spot syndrome virus. Indian Journal of Virology. 2012 Sep 14;23(2):161–74.

30. Components of the Immune System. In: Primer to the Immune Response. Elsevier; 2014. p. 21–54.

31. Bellinger DL, Millar BA, Perez S, Carter J, Wood C, ThyagaRajan S, et al. Innervation of lymphoid organs: Clinical implications. Clin Neurosci Res. 2006 Aug;6(1–2):3–33.

32. Hao W, Takano T, Guillemette J, Papillon J, Ren G, Cybulsky A V. Induction of Apoptosis by the Ste20-like Kinase SLK, a Germinal Center Kinase That Activates Apoptosis Signal-regulating Kinase and p38. Journal of Biological Chemistry. 2006 Feb;281(6):3075–84.

33. Cybulsky A V., Takano T, Guillemette J, Papillon J, Volpini RA, Di Battista JA. The Ste20-like kinase SLK promotes p53 transactivation and apoptosis. American Journal of Physiology-Renal Physiology. 2009 Oct;297(4):F971–80.

34. Bagaria J, Bagyinszky E, An SSA. Genetics of Autosomal Recessive Spastic Ataxia of Charlevoix-Saguenay (ARSACS) and Role of Sacsin in Neurodegeneration. Int J Mol Sci. 2022 Jan 4;23(1):552.

35. Vavassori S, Chou J, Faletti LE, Haunerdinger V, Opitz L, Joset P, et al. Multisystem inflammation and susceptibility to viral infections in human ZNFX1 deficiency. Journal of Allergy and Clinical Immunology. 2021 Aug;148(2):381–93.

36. Woodson CM, Kehn-Hall K. Examining the role of EGR1 during viral infections. Front Microbiol. 2022 Oct 21;13.

37. Browne EP, Wing B, Coleman D, Shenk T. Altered Cellular mRNA Levels in Human Cytomegalovirus-Infected Fibroblasts: Viral Block to the Accumulation of Antiviral mRNAs. J Virol. 2001 Dec 15;75(24):12319–30.

38. Shu M, Du T, Zhou G, Roizman B. Role of activating transcription factor 3 in the synthesis of latency-associated transcript and maintenance of herpes simplex virus 1 in latent state in ganglia. Proceedings of the National Academy of Sciences. 2015 Sep 29;112(39).

39. Rynes J, Donohoe CD, Frommolt P, Brodesser S, Jindra M, Uhlirova M. Activating Transcription Factor 3 Regulates Immune and Metabolic Homeostasis. Mol Cell Biol. 2012 Oct 1;32(19):3949–62.

40. Harwood SL, Nielsen NS, Diep K, Jensen KT, Nielsen PK, Yamamoto K, et al. Development of selective protease inhibitors via engineering of the bait region of human α2-macroglobulin. Journal of Biological Chemistry. 2021 Jul;297(1):100879.

41. Boons E, Vanstreels E, Jacquemyn M, Nogueira TC, Neggers JE, Vercruysse T, et al. Human Exportin-1 is a Target for Combined Therapy of HIV and AIDS Related Lymphoma. EBioMedicine. 2015 Sep;2(9):1102–13.

42. Uddin MdH, Zonder JA, Azmi AS. Exportin 1 inhibition as antiviral therapy. Drug Discov Today. 2020 Oct;25(10):1775–81.

43. Yarbrough ML, Mata MA, Sakthivel R, Fontoura BMA. Viral Subversion of Nucleocytoplasmic Trafficking. Traffic. 2014 Feb 2;15(2):127–40.

44. Azizian NG, Li Y. XPO1-dependent nuclear export as a target for cancer therapy. J Hematol Oncol. 2020 Dec 1;13(1):61.

45. Büssing I, Yang JS, Lai EC, Großhans H. The nuclear export receptor XPO-1 supports primary miRNA processing in C. elegans and Drosophila. EMBO J. 2010 Jun 2;29(11):1830–9.

46. Wolff T, O’Neill RE, Palese P. NS1-Binding Protein (NS1-BP): a Novel Human Protein That Interacts with the Influenza A Virus Nonstructural NS1 Protein Is Relocalized in the Nuclei of Infected Cells. J Virol. 1998 Sep;72(9):7170–80.

47. Zhang K, Shang G, Padavannil A, Wang J, Sakthivel R, Chen X, et al. Structural–functional interactions of NS1-BP protein with the splicing and mRNA export machineries for viral and host gene expression. Proceedings of the National Academy of Sciences. 2018 Dec 26;115(52).

48. Masson JJR, Cherry CL, Murphy NM, Sada-Ovalle I, Hussain T, Palchaudhuri R, et al. Polymorphism rs1385129 Within Glut1 Gene SLC2A1 Is Linked to Poor CD4+ T Cell Recovery in Antiretroviral-Treated HIV+ Individuals. Front Immunol. 2018 May 17;9.

49. Manel N, Kim FJ, Kinet S, Taylor N, Sitbon M, Battini JL. The Ubiquitous Glucose Transporter GLUT-1 Is a Receptor for HTLV. Cell. 2003 Nov;115(4):449–59.

50. Vittori A, Breda C, Repici M, Orth M, Roos RAC, Outeiro TF, et al. Copy-number variation of the neuronal glucose transporter gene SLC2A3 and age of onset in Huntington’s disease. Hum Mol Genet. 2014 Jun 15;23(12):3129–37.

51. Sun X, Xue C, Jin Y, Bian C, Zhou N, Sun S. Glucose transporter GLUT1 expression is important for oriental river prawn (Macrobrachium nipponense) hemocyte adaptation to hypoxic conditions. Journal of Biological Chemistry. 2023 Jan;299(1):102748.

52. Martínez-Quintana JA, Peregrino-Uriarte AB, Gollas-Galván T, Gómez-Jiménez S, Yepiz-Plascencia G. The glucose transporter 1 -GLUT1- from the white shrimp Litopenaeus vannamei is up-regulated during hypoxia. Mol Biol Rep. 2014 Dec 29;41(12):7885–98.

53. Wang XD, Li EC, Chen K, Wang SF, Li TY, Xu C, et al. Response of facilitative glucose transporter 1 to salinity stress and dietary carbohydrate nutrition in white shrimp *Litopenaeus vannamei*. Aquac Nutr. 2017 Feb;23(1):90–100.

54. Shi X, Meng X, Kong J, Luan S, Luo K, Cao B, et al. Transcriptome analysis of ‘Huanghai No. 2’ Fenneropenaeus chinensis response to WSSV using RNA-seq. Fish Shellfish Immunol. 2018 Apr;75:132–8.

55. Verbruggen B, Bickley L, Van Aerle R, Bateman K, Stentiford G, Santos E, et al. Molecular Mechanisms of White Spot Syndrome Virus Infection and Perspectives on Treatments. Viruses. 2016 Jan 18;8(1):23.

56. Millard RS, Bickley LK, Bateman KS, Farbos A, Minardi D, Moore K, et al. Global mRNA and miRNA Analysis Reveal Key Processes in the Initial Response to Infection with WSSV in the Pacific Whiteleg Shrimp. Viruses. 2021 Jun 13;13(6):1140.

57. Agnello V, Ábel G, Elfahal M, Knight GB, Zhang QX. Hepatitis C virus and other Flaviviridae viruses enter cells via low density lipoprotein receptor. Proceedings of the National Academy of Sciences. 1999 Oct 26;96(22):12766–71.

58. Nikolic J, Belot L, Raux H, Legrand P, Gaudin Y, A. Albertini A. Structural basis for the recognition of LDL-receptor family members by VSV glycoprotein. Nat Commun. 2018 Mar 12;9(1):1029.

59. Wang S, Li H, Weng S, Li C, He J. White Spot Syndrome Virus Establishes a Novel IE1/JNK/c-Jun Positive Feedback Loop to Drive Replication. iScience. 2020 Jan;23(1):100752.

60. Peppenelli MA, Miller MJ, Altman AM, Cojohari O, Chan GC. Aberrant regulation of the Akt signaling network by human cytomegalovirus allows for targeting of infected monocytes. Antiviral Res. 2018 Oct;158:13–24.

61. López-Cabrera M, Letovsky J, Hu KQ, Siddiqui A. Transcriptional factor C/EBP binds to and transactivates the enhancer element II of the hepatitis B virus. Virology. 1991 Aug;183(2):825–9.

62. Yang GJ, Wu J, Leung CH, Ma DL, Chen J. A review on the emerging roles of pyruvate kinase M2 in anti-leukemia therapy. Int J Biol Macromol. 2021 Dec;193:1499–506.

63. Lado S, Elbers JP, Plasil M, Loney T, Weidinger P, Camp J V., et al. Innate and Adaptive Immune Genes Associated with MERS-CoV Infection in Dromedaries. Cells. 2021 May 23;10(6):1291.

64. Javier RT, Rice AP. Emerging Theme: Cellular PDZ Proteins as Common Targets of Pathogenic Viruses. J Virol. 2011 Nov 15;85(22):11544–56.

65. Tanaka S, Tanaka K, Magnusson F, Chung Y, Martinez GJ, Wang Y hong, et al. CCAAT/Enhancer-Binding Protein α Negatively Regulates IFN-γ Expression in T Cells. The Journal of Immunology. 2014 Dec 15;193(12):6152–60.

66. Badu P, Baniulyte G, Sammons MA, Pager CT. Activation of ATF3 via the Integrated Stress Response Pathway Regulates Innate Immune Response to Restrict Zika Virus. 2023.

67. Arunachalam B, Phan UT, Geuze HJ, Cresswell P. Enzymatic reduction of disulfide bonds in lysosomes: Characterization of a Gamma-interferon-inducible lysosomal thiol reductase (GILT). Proceedings of the National Academy of Sciences. 2000 Jan 18;97(2):745–50.

68. Chen C, Zhang X, Lo C, Liu P, Liu Y, Gallo RL, et al. The essentiality of α-2-macroglobulin in human salivary innate immunity against new H1N1 swine origin influenza A virus. Proteomics. 2010 Jun 15;10(12):2396–401.

69. Jia X, Yamamura T, Gbadegesin R, McNulty MT, Song K, Nagano C, et al. Common risk variants in NPHS1 and TNFSF15 are associated with childhood steroid-sensitive nephrotic syndrome. Kidney Int. 2020 Nov;98(5):1308–22.

70. Shin SM, Moehring F, Itson-Zoske B, Fan F, Stucky CL, Hogan QH, et al. Piezo2 mechanosensitive ion channel is located to sensory neurons and nonneuronal cells in rat peripheral sensory pathway: implications in pain. Pain. 2021 Nov;162(11):2750–68.

71. Príncipe C, Dionísio de Sousa IJ, Prazeres H, Soares P, Lima RT. LRP1B: A Giant Lost in Cancer Translation. Pharmaceuticals. 2021 Aug 24;14(9):836.

